# A cytokinin-auxin antagonistic module regulates nitrogen-triggered tiller outgrowth in rice

**DOI:** 10.1101/2025.09.24.678274

**Authors:** Sourav Chatterjee, Aditi Dwivedi, Ananda K. Sarkar, Aashish Ranjan

## Abstract

Tillering is a key trait that shapes aboveground architecture and directly contributes to rice yield. However, integration of environmental cues to genetic regulators for controlling tiller outgrowth remains poorly understood. We investigated the effects of nitrogen, one of the critical environmental factors, on the early stages of tiller bud outgrowth. Comprehensive temporal transcriptomic analyses in response to two nitrogen forms, nitrate (NO_3_ ^-^) and ammonium (NH_4_ ^+^), identified gene-regulatory networks for nitrogen-triggered tiller outgrowth, with phytohormone signaling as the central interface. Pharmacological, molecular, and genetic experiments established cytokinin–auxin antagonism as a central regulator of the nitrogen-mediated tillering. Cytokinin promoted bud outgrowth by repressing the TCP transcription factor *OsTB1*, the key inhibitor of tillering, via Cytokinin Response Factors OsERF53/54. OsERF53/54 further reduced polar auxin transport by repressing *OsPIN1a* to promote bud activation. Conversely, auxin promoted bud dormancy by inducing *OsTB1* and *OsPIN1a* via OsARF11/16. Both NO_3_ ^-^ and NH_4_ ^+^ triggered tiller outgrowth and induced largely overlapping transcriptional responses, with a temporal delay for NO_3_ ^-^. Adequate nitrogen supplies induced cytokinin signaling while inhibiting auxin signaling and transport in tiller buds, thereby triggering bud outgrowth. Together, we identify a key regulatory role of auxin–cytokinin antagonistic module for integrating nitrogen signals to determine tiller bud fate.

## Introduction

In cereal grasses, tillers are specialized shoot branches that grow from axillary meristems at the lower leaf axils. These tillers may terminate in individual panicles upon subsequent reproductive transition. The number, position, angle, and growth of tillers significantly shape above-ground plant architecture and strongly influence cereal crop yield (Mathan *et al*., 2016). Tillers resemble the main culm soon after emergence in the early vegetative phase and often form adventitious roots, giving them partial independence from the primary culm. This sets them apart from dicot axillary branches, which emerge after reproductive transition, do not develop roots, and remain dependent on the main stem (Doust, 2007; Yang & Hwa, 2008; Kebrom *et al*., 2013). Despite the anatomical distinction, the key steps of tiller development in monocots and axillary branching in dicots share conserved mechanisms (Barbier *et al*., 2019; Luo *et al*., 2021). While genetic factors largely govern tiller bud establishment in the early phases of tiller development, a complex interplay of genetic regulators with physiological and environmental cues regulates the later outgrowth of buds, which is a key determinant of effective tiller number. The number of productive tillers is highly sensitive to environmental factors, often overriding genotypic potential, among all key architectural traits that affect the yield of rice, a major cereal crop (Takai, 2024). Therefore, a mechanistic understanding of the early stages of tiller bud outgrowth is crucial for optimizing rice yield potential in fluctuating environments.

In recent years, forward genetics has identified several molecular regulators of tiller bud outgrowth in rice. *OsTB1*, a TCP transcription factor orthologous to *ZmTB1* and *AtBRC1*, acts as a central repressor of this process (Takeda *et al*., 2003; Wang *et al*., 2018b). Studies using High Tillering/Dwarf (HTD) mutants identified genes involved in strigolactone (SL) biosynthesis and signaling as major suppressors of tiller outgrowth (Ishikawa *et al*., 2005). This SL-mediated inhibition of shoot branching is conserved across species (Wang *et al*., 2018b; Barbier *et al*., 2019). The SL-signaling repressor OsD53 inhibits *OsTB1* expression. Higher SL levels relieve this repression by degrading OsD53, subsequently inhibiting bud outgrowth (Song *et al*., 2017). Other known tillering regulators in rice also function by either modulating *OsTB1* expression or altering SL biosynthesis or signaling. More recently, OsBZR1, OsDLT, and OsRLA1, targets of brassinosteroid (BR) signaling, were found to interact with OsD53 and synergistically suppress *OsTB1* expression, integrating SL and BR signaling to antagonistically influence tiller outgrowth (Fang *et al*., 2020). Among other plant hormones, Gibberellins (GA) also restrict tiller outgrowth. GA signaling inhibitor, OsSLR1, promotes tiller outgrowth by preventing proteasomal degradation of OsMOC1 and OsNGR5. OsNGR5 consequently represses the SL receptor *OsD14*, further integrating GA signaling into the SL-OsTB1 module (Liao *et al*., 2019; Wu *et al*., 2020). Beyond SL, BR, and GA, genetic evidences indicate that other phytohormones also play critical roles in regulating rice tiller bud outgrowth. Mutants with impaired auxin signaling or transport, as well as those with disrupted cytokinin transport or metabolism, highlight the contributions of these hormones in tiller growth (Xia *et al*., 2012; Yeh *et al*., 2015; Jin *et al*., 2016; Zhao *et al*., 2019). However, integration of these phytohormones with regulatory pathways known to be involved in tillering remains elusive. A comprehensive understanding of tiller bud outgrowth requires integration of these molecular components within the broader regulatory networks that coordinate this process.

Phytohormones simultaneously sense and integrate fluctuating environmental signals into the regulatory network of tiller development while governing the tiller outgrowth decision. Although several environmental cues affect tiller outgrowth, soil nutrient content remains the limiting factor for major cultivated crops like rice. Among nutrients, nitrogen significantly influences tillering and has been shown to trigger the outgrowth of dormant tiller buds (Luo *et al*., 2017). The introduction of chemical nitrogen fertilizers alongside semi-dwarf and high-tillering Green Revolution rice varieties (GRVs) has dramatically increased rice yields over the past five decades (Luo *et al*., 2020). Genetic studies have identified transcription factors, such as *OsNGR5*, *OsTCP19*, and *OsGATA8*, as key regulators of nitrogen-triggered tillering in GRVs (Wu *et al*., 2020, 2024; Liu *et al*., 2021). However, yield gains in GRVs have plateaued in the last few decades due to their low nitrogen use efficiency (NUE) (Asami, 2023). Recent findings highlight that DELLA-mediated inhibition of OsGRF4 causes reduced NUE in GRVs, with OsGRF4 integrating gibberellin and strigolactone signaling pathways to balance tiller outgrowth and NUE (Li *et al*., 2018; Sun *et al*., 2023). These advancements imply shared phytohormonal regulation of tiller outgrowth and nitrogen-responsive signaling associated with NUE. Additionally, rice shows different responses to the two major nitrogen forms, nitrate (NO₃⁻) and ammonium (NH₄⁺), resulting in distinct tillering patterns (Yuan Wang *et al*., 1993; Yi *et al*., 2019b). A recent study reported that *OsPIN9* downstream of these nitrogen forms contributes to this difference, inferring that auxin transport likely plays a critical role in mediating distinct responses to different nitrogen forms (Hou *et al*., 2021a). However, a more comprehensive investigation is necessary to elucidate the integration of phytohormone signaling with nitrogen sensing to trigger the transition from bud dormancy to active tiller growth.

Although the genetic regulation of tillering in rice has been investigated, the integration of environmental cues with these genetic components is rather fragmented. Particularly, limited attempts have been made to connect nitrogen supply with genetic regulators into a coherent framework during the early transition from dormant bud to active tiller outgrowth. In this study, we aimed to fill this gap by investigating the effects of nitrogen availability on global gene expression and underlying genetic regulation during the early stages of nitrogen-triggered bud outgrowth. Our results confirm the central role of phytohormone signaling in nitrogen-triggered tiller development. We further highlight the role of auxin–cytokinin antagonism, a regulatory layer not yet explored in rice tillering. We show that cytokinin and auxin converge on *OsTB1* and *OsPIN1a*, with opposing effects on their transcription. Nitrogen supply shifts this balance by enhancing cytokinin signaling and restricting auxin signaling and transport in tiller buds, leading to bud activation.

## Materials and methods

### Plant materials and growth conditions

Most experiments were conducted with *Oryza sativa* ssp. *Indica* cv. IR64 seedlings, except for the natural variation studies. Rice accessions used for the natural variation studies are taken from the IRRI 3k panel (Wang *et al*., 2018a) and are listed in Supplementary Table S1. Rice seeds were surface sterilized with 30% bleach followed by 200 mg/L Bavistin, then germinated on wet germination paper, followed by their transfer to hydroponic culture with modified Yoshida medium (Yoshida *et al*., 1976; Sharma *et al*., 2018). Medium composition is provided in Supplementary Table S2. The nutrient solution was replaced every third day during seedling growth. Plants were grown at 28 °C day/25 °C night, 16 h light/8 h dark, 230–250 μmol m⁻² s⁻¹ white light, and 70% relative humidity. Tobacco (*Nicotiana benthamiana*) seedlings were grown in a 1:1 mixture of nutrient soil and vermiculite at 25 °C day/22 °C night, under 16 h light/8 h dark at 120–150 μmol m⁻² s⁻¹ cool white light.

### RNA extractions, qPCR, RNA-seq library preparation, and sequencing

For expression analyses, rice seedlings were grown until the 5th leaves were fully expanded and the 6th leaves had just emerged. Seedlings were then transferred to specific treatment media for defined durations. After said duration, tiller buds from the fourth leaf axils were dissected under a stereo microscope for RNA extraction. For expression assays with cycloheximide (CHX), seedlings were pre-incubated in minimal nitrogen media containing 25 μM CHX or DMSO, and then transferred for 6 h to either minimal nitrogen media with CHX or DMSO, or to 1 μM kinetin-supplemented media with CHX or DMSO. All other hormone treatments for expression analysis were also performed for 6 h unless stated otherwise. For qPCR, Oligo(dT)-primed cDNA was synthesized using the Verso cDNA Synthesis Kit (Thermo Fisher Scientific). qPCR reactions were run on an ABI QuantStudio 6 Flex with gene-specific primers and PowerUp™ SYBR™ Green master mix (Applied Biosystems). Primer sequences are listed in Supplementary Table S3. cDNA libraries for RNA-seq were prepared in three biological replicates using the YourSeq Full Transcript RNA-seq Library Kit (Amaryllis Nucleics, Oakland, CA, USA) following the manufacturer’s instructions, and sequenced on the Illumina HiSeq 4000 platform.

### RNA-seq analysis and network construction

Quality-filtered reads were mapped to representative Oryza sativa cDNA sequences from the Rice Genome Annotation Project (RGAP7) database using default parameters of Bowtie2 (Langmead & Salzberg, 2012; Hamilton *et al*., 2025). Differential expression was analyzed by doing pairwise comparisons with the edgeR package (Robinson & Oshlack, 2010) using a quasi-likelihood F-test. Significantly differentially expressed genes (DEGs) in each pair-wise comparison were identified based on FDR < 0.05 and logFC ≥ 0.585 or ≤ –0.585 (Supplementary dataset S1). GO enrichment was performed using the tools from PlantRegMap of the Plant Transcription factor database (Tian *et al*., 2020), considering biological process terms with q < 0.05 (Supplementary dataset S2).

Weighted Gene Co-expression Network Analysis (WGCNA) (Zhang & Horvath, 2005; Langfelder & Horvath, 2008) was performed using the top 80% variance DEGs. A soft-threshold power of 9 (R² ≥ 0.8) was used to generate a signed weighted adjacency matrix, transformed into a topological overlap matrix (TOM) using signed hybrid TOMtype. Genes were hierarchically clustered, and co-expression modules were identified with cutreeDynamic (minModuleSize = 30, deepSplit = 2). Similar modules were merged at mergeCutHeight = 0.15. Genes having MM ≥ 0.8 and MMpvalue < 0.01 were defined as Hub genes for said module (Supplementary dataset S3).

Module-wise co-expression networks were built from hub genes of interest (GOIs) with max_neighbors = 50 and tom_cutoff = 0.15. Pearson correlation coefficients (PCCs) between intermodular GOIs were represented as colored edges to connect these co-expression networks. For Transcription factor (TF)-target networks, genomic sequences (–2 kb to +1 kb from transcription start site/ TSS) of targets and TFs were scanned for TF-binding sites using FIMO (MEME-suite_5.5.8) with motifs from the Cis-BP_2.00 database (Grant *et al*., 2011; Weirauch *et al*., 2014). Regulator-target pairs were inferred from expression data using GENIE3 (Huynh-Thu *et al*., 2010), and only FIMO-supported pairs were retained in the networks. Nodes and edges were collapsed by gene family and module assignment (e.g., type-B response regulators/BRRs in blue module represented as one node) for the gene regulatory network from the turquoise and blue modules

### EdU labelling and visualization

Steps of EdU (5-ethynyl-20-deoxyuridine) labelling are shown schematically in Fig. S1A. Shoot bases with the fourth axil tiller bud were dissected after 12 hours of treatment in a particular medium, and then incubated in the same medium containing 10 µM EdU at 28 °C, 230–250 μmol m⁻² s⁻¹ white light, and 70% relative humidity. After 1 h (pulse), samples were washed and re-incubated in similar conditions without EdU (chase) (Yin & Tsukaya, 2016). Following chase, shoot bases were fixed in chilled 4% paraformaldehyde in PBS, dehydrated through successive ethanol-PBS and xylene-ethanol gradients, and then embedded in paraffin. Longitudinal sections (10 µm) were cut from the embedded tissue through tiller buds. Sections were then dewaxed, rehydrated, and EdU-labelled cells were detected with the Click-iT™ EdU Cell Proliferation Kit (Invitrogen). Sections were counterstained with Calcofluor White stain (Fluka #18909). Images were acquired on a Leica TCS SP8 confocal microscope (10× objective, Alexa Fluor 488 settings: 488 nm excitation, 500-540 nm emission; Calcofluor White: 360 nm excitation, 430 nm emission).

### Yeast two-hybrid (Y2H) assays

Y2H assays used the GAL4-based two-hybrid system (Clontech, Mountain View, CA, USA). Full-length coding sequences of OsERFs and OHPs were cloned into pENTR-D-TOPO (Invitrogen #K240020). OHP1/2-AD prey constructs were generated via LR reactions into pGADT7-GW, and OsERF53/54-BD bait constructs into pGBKT7-GW. OsRR24-BD served as a positive control (Zhao *et al*., 2018). Yeast transformations (strain Y2H Gold; Takara #630489) were performed using the EZ yeast transformation kit (MP Biomedicals) following the manufacturer’s protocol. Interactions were assessed by co-transforming each prey construct with each bait construct or an empty pGBKT7-GW. An empty pGADT7-GW was co-transformed with OsERF-BD bait constructs as negative controls.

### Dual-luciferase assays

Promoter–reporter constructs included two *OsTB1* promoter fragments (–1 to –1991 bp; –1474 bp to +930 bp from the start codon) and the *OsPIN1a* promoter (–1 to –2000 bp from the start codon), cloned upstream of Firefly luciferase in pGreenII 0800-LUC binary vector, having CaMV35S-driven Renilla Luciferase as an internal control. OsERF53/54 and OsARF11/16 CDSs were cloned into pENTR-D-TOPO and transferred into pEarlyGate101 to generate TF-YFP fusions; YFP-expressing empty binary vector pEarlyGate104 was used as a negative control. Promoter constructs were co-transformed with pSoup into Agrobacterium GV3101, and co-infiltrated into *N. benthamiana* leaves with TF or control constructs. After 48 hours post-infiltration, leaves were sprayed with 150ng/μl D-luciferin potassium salt (GoldBio) and imaged with a ChemiDoc MP imaging system (Bio-Rad). For dual-luciferase assays, leaf discs of 1 cm diameter were collected, and Firefly/Renilla luciferase activities were measured using the Dual-Luciferase Assay System (Promega) and POLARstar Omega (BMG Labtech).

### Quantification, visualization, and statistical analysis

Tiller buds up to < 2 mm were imaged under a stereomicroscope, and lengths were measured in ImageJ. Buds > 2 mm were measured with a hand ruler. EdU fluorescence was also quantified using ImageJ. qRT–PCR data were plotted in GraphPad Prism 10. Other datasets were visualized in RStudio (packages listed in Supplementary Table S4). Gene regulatory networks were visualized using Cytoscape (Shannon *et al*., 2003). Statistical analyses for RNA-seq were performed within the respective pipelines. Other data were analyzed using one-way/two-way ANOVAs with post-hoc tests (GraphPad Prism 10) or two-tailed Student’s t-tests (Microsoft Excel). A two-tailed t-test for the correlation of ERF expression and phenotype was also done in Microsoft Excel. Fisher’s exact tests for DEG overlaps were done in a web-based tool (https://www.socscistatistics.com/tests/fisher/default2.aspx).

## Results

### Exogenous nitrogen supply promotes rice tiller bud outgrowth

To investigate the role of nitrogen (N) during the early stages of rice tiller bud outgrowth, we estimated the effects of exogenous application of two different N forms, ammonium (NH_4_^+^/Am) and nitrate (NO_3_^-^/Ni), on the early bud growth. *Oryza sativa* ssp. *Indica* cv. IR64 seedlings were grown on minimal N, which was adequate to sustain seedling growth but insufficient to promote tiller outgrowth. Seedlings were then transferred to media supplemented with sufficient N, either as ammonium (NH_4_^+^/Am) or nitrate (NO_3_^-^/Ni), once the 5th leaves were fully expanded (Fig. 1A). Both ammonium and nitrate supply significantly increased the rice tiller bud outgrowth in the 4th leaf axis. While ammonium treatment increased tiller bud length significantly at 3 days after transfer (DAT), nitrate-supplied plants showed significant tiller bud elongation by 4 DAT (Fig. 1B, C). Furthermore, ammonium treatment had a more pronounced effect on tiller bud outgrowth, as tiller buds in ammonium-treated seedlings were consistently longer than in nitrate-treated ones. To explore cellular changes underlying N-mediated tiller bud outgrowth, we performed an EdU-based cell proliferation assay (Fig. S1A). Within 12 hours of sufficient N supply, tiller buds exposed to either ammonium or nitrate exhibited a higher frequency of EdU-stained proliferating cells compared to those under nitrogen-deficient conditions. However, tiller buds in nitrate-treated seedlings displayed fewer EdU-stained cells than ammonium-treated ones, aligning with the slower phenotypic response of seedlings to nitrate treatment (Fig. 1D, E). We also analyzed the expression of genes related to cell division and hormone signaling and metabolism, which are known to affect tiller outgrowth in rice, at early time points following ammonium or nitrate supplementation (Fig. S1B). The mitosis marker gene *OsCDKB2* showed early induction, with peak expression at 12 hours post-transfer (Fig. S1C). Most hormone-related genes also responded within 2 – 12 hours of nitrogen supplementation (Fig. S1C), suggesting that the transcriptional switch from dormancy to active growth occurs within this window. Taken together, our results demonstrated that sufficient nitrogen supply induces the outgrowth of dormant tiller buds via enhancing cell division within 12 hours, marking the transition to active growth. Moreover, the superior effect of ammonium in promoting tiller outgrowth was evident at both phenotypic and cellular levels.

**Fig. 1.**
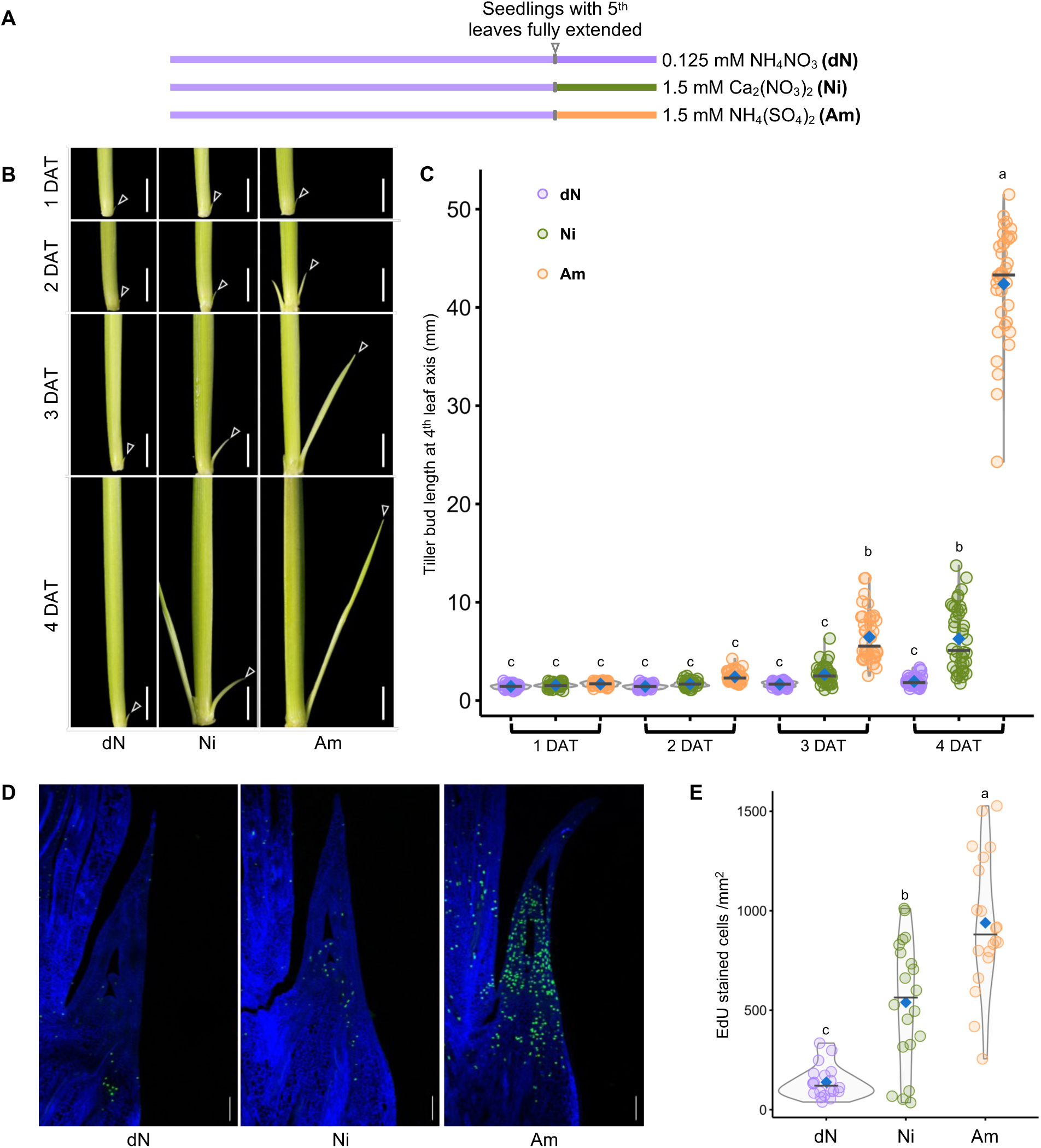
Exogenous nitrogen supply promotes rice tiller bud outgrowth. (A) Schematic diagram showing the experimental conditions. Seedlings grown on minimal nitrogen-containing Yoshida media were supplemented with sufficient nitrate (Ni) or ammonium-containing media (Am). Control seedlings were continued to grow in minimal nitrogen media (dN). Nitrogen salts and concentrations are mentioned in the figure. (B) & (C) Representative images (B, Scale = 1cm) and quantification (C) of tiller bud length at the 4^th^ leaf axil (marked by white triangles in B) of seedlings over four days post-supplementation. The blue diamonds and the horizontal lines in (C) represent mean and median values, respectively (n=40). The lowercase letters indicate significant differences based on two-way ANOVA and Tukey’s test (*p* < 0.05). (D) & (E) Representative images (D, Scale = 100 µm) and quantification (E) of EdU-stained cells in tiller buds at the 4^th^ leaf axil at 12 hours after supplementation. The blue diamonds and the horizontal lines in (E) represent mean and median values, respectively (n=15). The lowercase letters indicate significant differences based on one-way ANOVA and Tukey’s test (*p* < 0.05).

### Transcriptome profiling identified auxin and cytokinin as the key regulators of nitrogen-mediated tiller bud outgrowth

To explore the genetic mechanisms underlying the nitrogen-mediated tiller bud outgrowth, we compared the transcriptome of the tiller buds at the 4th leaf axis grown under minimal nitrogen (dN) and at 2– and 12-hours after supplementing with either sufficient ammonium (Am) or nitrate (Ni). Clustering of samples based on gene expression grouped Ni-supplied samples at 2 hours with dN samples at both time points, suggesting minimal early transcriptional changes in response to nitrate (Fig. 2A). In contrast, ammonium triggered distinct gene expression changes at 2 hours post-supplementation. Interestingly, both ammonium– and nitrate-treated samples exhibited overlapping gene-expression patterns at 12 hours, which differed markedly from those of the dN samples (Fig. 2A). Consistent with this pattern, nitrate-supplied buds showed only 63 differentially expressed genes (DEGs, p < 0.05, |fold change| >1.5, Supplementary dataset S1) at 2 hours relative to dN, 28 upregulated and 35 downregulated (Fig. 2B). Despite limited number of DEGs, GO terms such as “nitrate metabolic process,” “nitrate assimilation,” and “ammonium ion metabolism” were significantly enriched in upregulated genes (Fig. S2A, Supplementary dataset S2). In contrast, ammonium induced 2,087 and suppressed 1,048 genes at 2 hours. While upregulated genes were significantly enriched in GO terms like “cell cycle,” “chromosome organization,” “response to cytokinin,” and “amino acid metabolic process,” downregulated genes were associated with “response to starvation,” “amino acid catabolic process,” and terms related to phytohormone signaling and response (Fig. S2A, B).

**Fig. 2.**
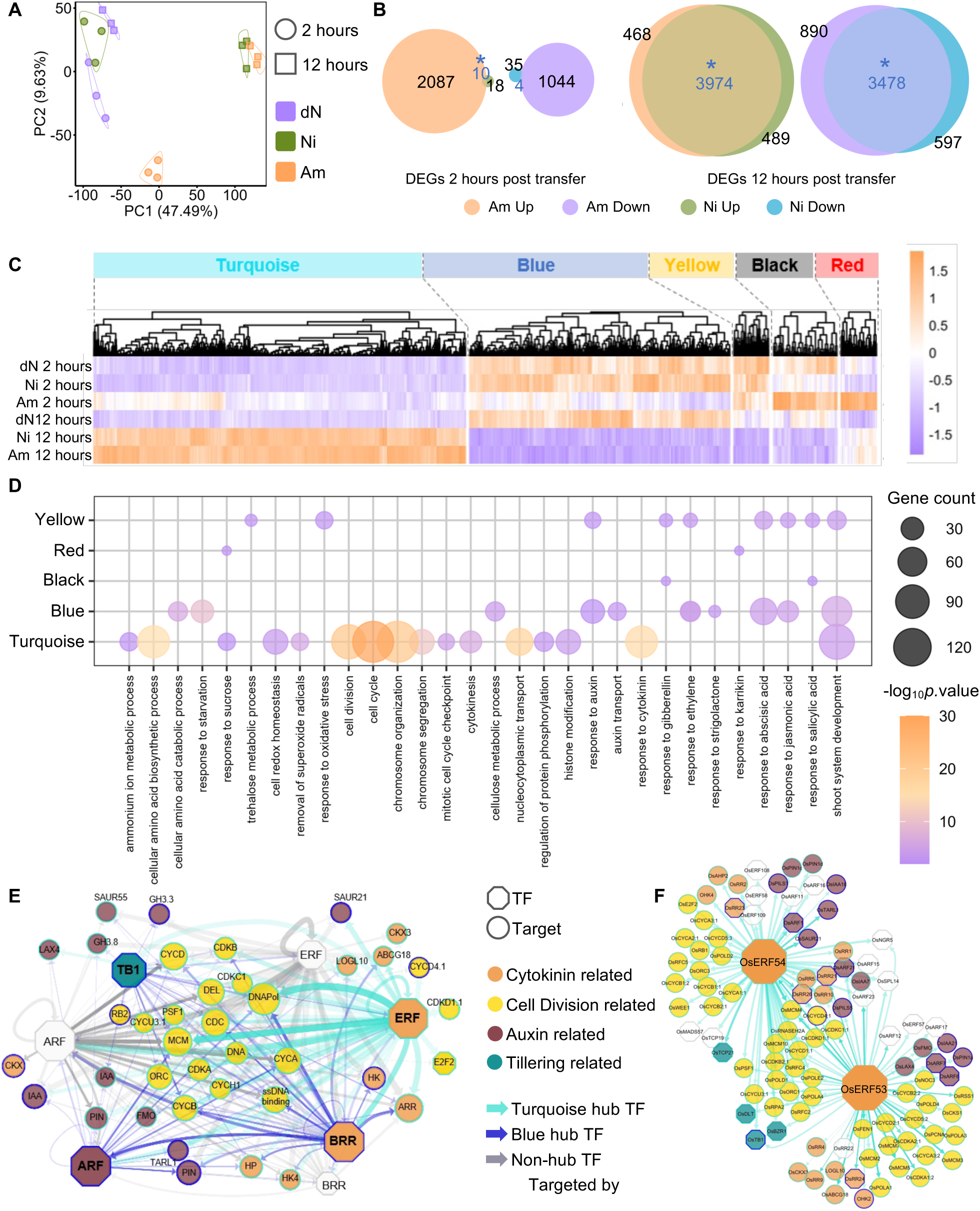
Transcriptome profiling reveals auxin and cytokinin as the key regulators of nitrogen-mediated tiller bud outgrowth. (A) Principal-component analysis (PCA) of gene-expression profile in tiller buds from 4^th^ leaf axil of seedlings supplemented with minimal nitrogen/dN, sufficient nitrate/Ni, and sufficient ammonium/Am containing media, at 2– and 12-hours post-supplementation (n = 3 biological replicates). (B) Euler diagrams illustrating the numbers and overlaps of Up– and Downregulated DEGs (FDR<0.05 and |fold change| >1.5) in Am and Ni supplementation in comparison to dN. Blue numbers indicate the number of overlapping genes. Stars (*) on intersections represent significant overlap based on Fisher’s exact test (*p*<0.05). (C) Heatmaps representing the normalized expression values (z-score) of Hub genes from different WGCNA modules for each sample in a row. (D) Bubble plot displaying representative GO terms (biological process) enriched in the hub genes of each module. (E) Regulatory network for cell cycle, auxin, cytokinin, and tillering-related genes from the turquoise and blue modules. TFs and targets are collapsed according to family and module membership. Colors of node borders depict module assignment, and edge colors represent the module of regulatory TF. Collapsed edge thickness represents log(total binding sites), and edge transparency represents total edge weight. Unfilled TF nodes represent TFs from auxin or cytokinin pathways that are not hub-TFs from the blue or turquoise modules. (F) Regulatory network for cell cycle, auxin, cytokinin, and tillering-related genes from the turquoise and blue modules targeted by OsERF53 and OsERF54. Colors of node borders depict module assignment, and edge colors represent the module of regulatory TF. Edge thickness represents the number of binding sites from −2kb to +1kb from the TSS of target genes, and edge transparency represents edge weight. Unfilled TF nodes represent TFs from auxin or cytokinin pathways that are not hub-TFs from the blue or turquoise modules.

Both ammonium and nitrate-supplied buds showed remarkably higher numbers of DEGs compared to dN samples at 12 hours. Ammonium supplementation induced 4,442 genes and suppressed 4,368 genes, while nitrate supplementation induced 4,463 genes and suppressed 4,075 genes. A significant overlap of DEGs occurred, with 3,974 upregulated and 3,478 downregulated genes shared between both treatments (Fig. 2B). Significantly enriched GO terms that were common for upregulated genes at 12 hours of the two treatments included “response to cytokinin”, “chromosome segregation,” “cytokinesis,” and terms linked to sucrose and superoxide metabolism (Fig. S2A). Downregulated genes were significantly enriched for response to various stimuli and terms related to different hormone pathways (Fig. S2B). Together, these results showed that while both N forms eventually elicit similar transcriptomic changes, ammonium triggers earlier transcriptional reprogramming that facilitates faster tiller bud activation. This data further suggested strong involvement of hormone signaling and response in nitrogen-mediated tiller bud outgrowth.

To identify groups of genes with similar expression patterns and link them to different nitrogen-temporal regimes, we performed WGCNA on the top 80% most variable DEGs. Seven co-expression modules were detected. The turquoise (4,190 genes) and blue (3,367 genes) modules together comprised ∼91.7% of all genes (Fig. S3A, Supplementary dataset S3), but neither showed significant correlation (|correlation coefficient| > 0.7) with any individual sample. Instead, the red module (157 genes) was significantly correlated with 2 hours of ammonium supplementation, and the pink module (51 genes) with 12 hours of dN (Fig. S3B). Given that our samples displayed overlapping transcriptomic signatures in PCA (Fig. 2A), we performed hierarchical clustering on PCA coordinates, grouping samples into three distinct clusters (Fig. S3C, D). Correlating WGCNA modules to these clusters, rather than with individual samples, revealed a robust association pattern (Fig. S3E). The turquoise module, with the highest number of genes, was significantly associated with cluster 3 (12 hours after ammonium or nitrate supplementation), and negatively correlated with cluster 1 (minimal nitrogen and 2 hours nitrate supplementation). The blue module exhibited an opposite trend, with positive correlation with cluster 1 and negative correlation with cluster 3. The yellow (342 genes) and black (88 genes) modules also showed significant negative correlation with cluster 3 (Fig. S3E). We identified hub genes for each module. The pink and magenta modules had very few hub genes, 7 and 9, respectively, and showed no significant association with any sample clusters (Fig. S3A, E), and thus were excluded from further analysis. We then visualized expression of hub genes across samples, which reflected their cluster associations (Fig. 2C). Hub genes in the turquoise module were enriched in processes related to amino acid and sugar metabolism, cell division, and cytokinin signaling. The blue module hub genes were enriched in starvation related terms, and hormone pathways such as auxin, ABA, JA, ethylene, and strigolactone, consistent with its association with dormant tiller buds (Fig. 2D). Enriched pathways for the yellow module hub genes overlapped with the blue module in phytohormone pathways, in addition to showing enrichment of gibberellin related terms (Fig. 2D). Taken together, phytohormone pathways emerged as central regulators of dormancy release in response to nitrogen treatment, together with active metabolism and cell division.

To further explore the involvement of phytohormones in nitrogen-mediated tiller bud outgrowth, we curated canonical genes for phytohormone metabolism, transport, and signaling from the literature (Supplementary dataset S4), along with genes for cell-cycle, DNA replication, and transcription factors known to regulate rice tiller bud outgrowth. Canonical genes that were expressed across samples used for the transcriptomic comparisons and were hub genes in any of the detected modules were identified as genes of interest (GOIs) (Fig. S4A). We constructed a module-wise co-expression network of hub GOIs and their immediate neighbors, and visualized their correlation across modules (Fig. S4B). GOIs of the turquoise module were negatively correlated with those of the blue, yellow, and black modules, reflecting their contrasting associations with these clusters. The blue and turquoise modules consisted of most GOIs, which were highly connected within modules. Hence, we focused further analysis on these two antagonistic modules. Among phytohormone-related GOIs, hub genes related to auxin and cytokinin signaling and transport were most abundant (Fig. S4A). While “auxin-transport” and “response to auxin” were enriched in the blue module, “response to cytokinin” was enriched in the turquoise module (Fig. 2D). The opposing enrichment of auxin– and cytokinin-related terms in the two inversely correlated modules, together with the abundance of hub genes from these pathways, suggested a possible auxin–cytokinin antagonism. This antagonism could play a central role in nitrogen-triggered bud outgrowth.

Although genetic evidences suggest auxin and cytokinin to be involved in tiller bud outgrowth (Xia *et al*., 2012; Kamada-Nobusada *et al*., 2013; Zhao *et al*., 2019), their molecular integration with the existing tiller bud outgrowth networks remains ambiguous. Hence, we investigated the gene-regulatory networks for the influence of auxin and cytokinin on tiller growth and active cell division during the outgrowth process. To this end, 72 cell-division, 20 auxin-related, 23 cytokinin-related, and 5 tillering-related hub genes from the turquoise or blue modules were examined for potential regulation and cis-binding by auxin-or cytokinin-related transcription factors (TFs) from these modules. Because hormone signaling can regulate TFs at both transcriptional and post-translational levels, we also included 9 ARFs and 5 cytokinin-related TFs expressed in the samples that were not hub genes in these modules. We built a regulatory network using expression data and predicted TF binding sites on target genes, collapsing nodes and edges by gene family and module assignment (Fig. 2E). Surprisingly, BRRs, canonical cytokinin-related TFs, were in the blue module, linked to dormant tiller buds. Four ARFs were also predictably detected as hub TFs in the blue module, along with TCP transcription factor OsTB1, a central repressor of bud outgrowth (Minakuchi *et al*., 2010). These TFs were predicted to regulate multiple cell division, auxin, and cytokinin-related genes (Fig. 2E). Nine other ARFs, which were not detected as hub-TFs in either of the modules, also appeared as potential regulators. Subnetworks of these TFs and their targets highlighted their role in the auxin– cytokinin–cell division nexus during bud outgrowth (Fig. S5). However, type-A response regulators (ARRs), which are direct cytokinin targets (Tsai *et al*., 2012), were highly represented in the turquoise module, indicating strong cytokinin signaling activity within this co-expression network. Interestingly, Cytokinin Response Factors (CRF) *OsERF53* and *OsERF54* (Rashotte *et al*., 2006; Lei *et al*., 2024) emerged as only cytokinin-related hub TFs in the turquoise module (Fig. 2E), suggesting they may act as key drivers of nitrogen-induced bud outgrowth. Direct targets of these ERFs included several cell division and cytokinin-related genes, and especially, auxin-related genes and *OsTB1*, suggesting these ERFs as potential mediators of auxin-cytokinin crosstalk regulating tiller bud outgrowth (Fig. 2F).

### Cytokinin triggers tiller bud growth downstream of nitrogen supply

Since transcriptomic analysis indicated a key role of cytokinin in nitrogen-mediated transition from dormant to active tiller bud growth, we examined the expression of cytokinin signaling, transport, and metabolism genes across nitrogen regimes and time points (Fig. 3A). Several cytokinin signaling and response genes were differentially expressed in response to exogenous nitrogen. Notably, *ARR*s were upregulated in ammonium– and nitrate-treated tiller buds at 12 hours, supporting the involvement of active cytokinin signaling in nitrogen-mediated bud outgrowth. To confirm this, we tested the effect of kinetin, a synthetic adenine-type cytokinin, on bud outgrowth under minimal nitrogen. Kinetin supplementation promoted tiller bud outgrowth and mimicked the phenotype seen under sufficient nitrogen (Fig. 3B, C). This kinetin-induced outgrowth involved increased cell proliferation, as shown by more cells entering the S-phase within 12 hours of kinetin-supplementation, similar to nitrogen (Fig. 3D, E). These results suggest that cytokinin acts downstream in the nitrogen-triggered bud outgrowth pathway.

**Fig. 3.**
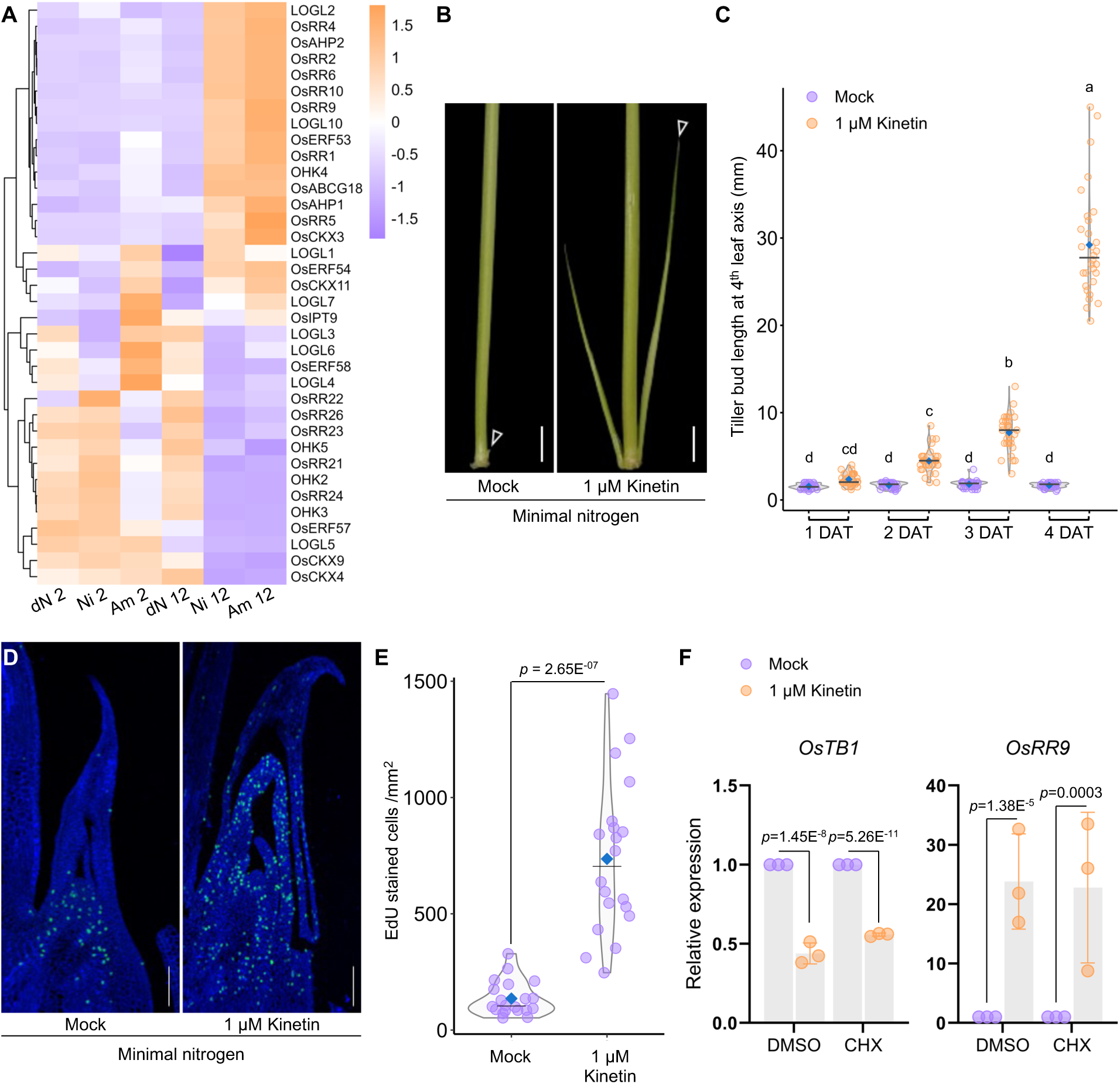
Cytokinin triggers tiller bud growth downstream of nitrogen supply. (A) Heatmap representing the normalized expression values (z-score) of canonical cytokinin-pathway genes in tiller buds at 2– and 12-hours across nitrogen regimes (dN/minimal nitrogen, Ni/sufficient nitrate, Am/sufficient ammonium). (B) Representative images of tiller bud length at the 4^th^ leaf axil (marked by white triangles) of seedlings grown in minimal nitrogen-containing media supplemented with 1μM Kinetin at four days post-supplementation. Scale = 1cm. (C) Quantification of tiller bud length at the 4^th^ leaf axil over 4 days post-supplementation. The blue diamonds and the horizontal lines represent mean and median values, respectively (n=30), different letters indicate significant differences based on two-way ANOVA and Tukey’s test, *p* < 0.05. (D) & (E) Representative images (D, Scale = 100 µm) and quantification (E) of EdU-stained cells in tiller buds at the 4^th^ leaf axil 12 hours after 1μM Kinetin supplementation in dN media. In E, the blue diamonds and the horizontal lines represent mean and median values, respectively (n=15, pairwise Student’s t test, *p*-value is indicated in the graph). (F) Relative expression of *OsTB1* and *OsRR9* in minimal nitrogen media with and without 1μM Kinetin, either in mock (DMSO) or in 25 μM Cycloheximide (CHX) supplementation (n=3; *p*-values obtained from pairwise Student’s t test are indicated in the graphs). *OsGAPDH* was used as the endogenous control.

*OsTB1*, the central repressor of tiller bud outgrowth, was identified as a hub-TF in the blue module and showed a negative correlation to the turquoise module enriched in active cytokinin signaling (Fig. 2E). Therefore, we examined whether cytokinin signaling integrates with *OsTB1* for enabling the shift from dormancy to active tiller growth under minimal nitrogen. Kinetin supplementation in a minimal nitrogen background significantly repressed the expression of *OsTB1* (Fig. 3F). In contrast, the treatment induced the expression of *OsRR9*, an *ARR* used as a positive control for a canonical cytokinin target (Tsai *et al*., 2012). To test if the kinetin-mediated repression of *OsTB1* is mediated by canonical cytokinin signaling, we quantified *OsTB1* expression in the presence of cycloheximide (CHX), a translational inhibitor. Kinetin-mediated *OsTB1* repression persisted with CHX treatment. *OsRR9* also maintained its expression pattern in CHX-treated samples. Taken together, these results indicate canonical cytokinin signaling suppresses *OsTB1* downstream of nitrogen supply to induce tiller bud outgrowth.

### OsERF53/54 suppress *OsTB1* expression downstream of cytokinin

Since *OsTB1* was suppressed by canonical cytokinin signaling, we first checked if BRRs could directly regulate *OsTB1*. Gene regulatory networks showed that BRRs did not directly target *OsTB1* and their expression was not associated with nitrogen supply or active bud growth (Fig. 3A, S5A). Instead, two hub-TFs of the turquoise module, OsERF53 and OsERF54, were found to directly regulate *OsTB1* (Fig. 2F). The expression of these two *ERFs* was strongly correlated with nitrogen supply and active tiller outgrowth (Fig. 3A). OsERF53/54 are members of a cytokinin-inducible subset of AP2-ERF transcription factors, known as CRFs, that act as a branch point in the cytokinin two-component phosphorelay system (Rashotte *et al*., 2006). Analyzing rice CRF expression (Lei *et al*., 2024) showed that only *OsERF53* and *OsERF54* were induced by sufficient nitrogen, with significant induction detectable within 2 hours of ammonium supplementation (Fig. S6A). Kinetin supplementation in minimal nitrogen media also significantly upregulated *OsERF53/54* (Fig. 4A). Both ERF proteins interacted with cytokinin phosphorelay components OHP1 and OHP2 in a yeast two-hybrid assay, further attesting their association with canonical two-component phosphorelay (Fig. 4B). These results suggested that OsERF53/54 may suppress *OsTB1* downstream of cytokinin signaling. We next searched for putative AP2/ERF binding sites, GCC box motifs, in the *OsTB1* gene and its upstream sequence (Fujimoto *et al*., 2000; Weirauch *et al*., 2014). One such GCC box was identified in the 5′ UTR of the *OsTB1* (Fig. S6B). To test the interaction with ERFs, we cloned the *OsTB1* promoter (*proOsTB1:fLuc*; –1991 bp upstream of the start codon) for transient dual-luciferase assays (Fig. S6C). Both OsERF53 and OsERF54 suppressed the *OsTB1* promoter activity (Fig. 4C, D). We also found several GCC box motifs within the *OsTB1* exon (Fig. S6B). To test whether OsERFs interact with these sites, we generated a translational fusion construct with a shorter *OsTB1* promoter driving firefly luciferase fused with exon sequences (*proOsTB1:OsTB1-fLuc*; −1474 bp upstream to +930 bp downstream of the start codon) (Fig. S6C). OsERF53 and OsERF54 also suppressed the activity of the translational fusion construct (Fig. 4E, F), confirming that they bind multiple GCC boxes within the *OsTB1* promoter and gene body to lower its expression. Together, these results show that canonical cytokinin signaling represses *OsTB1* through OsERF53/54 downstream of nitrogen supply.

**Fig. 4.**
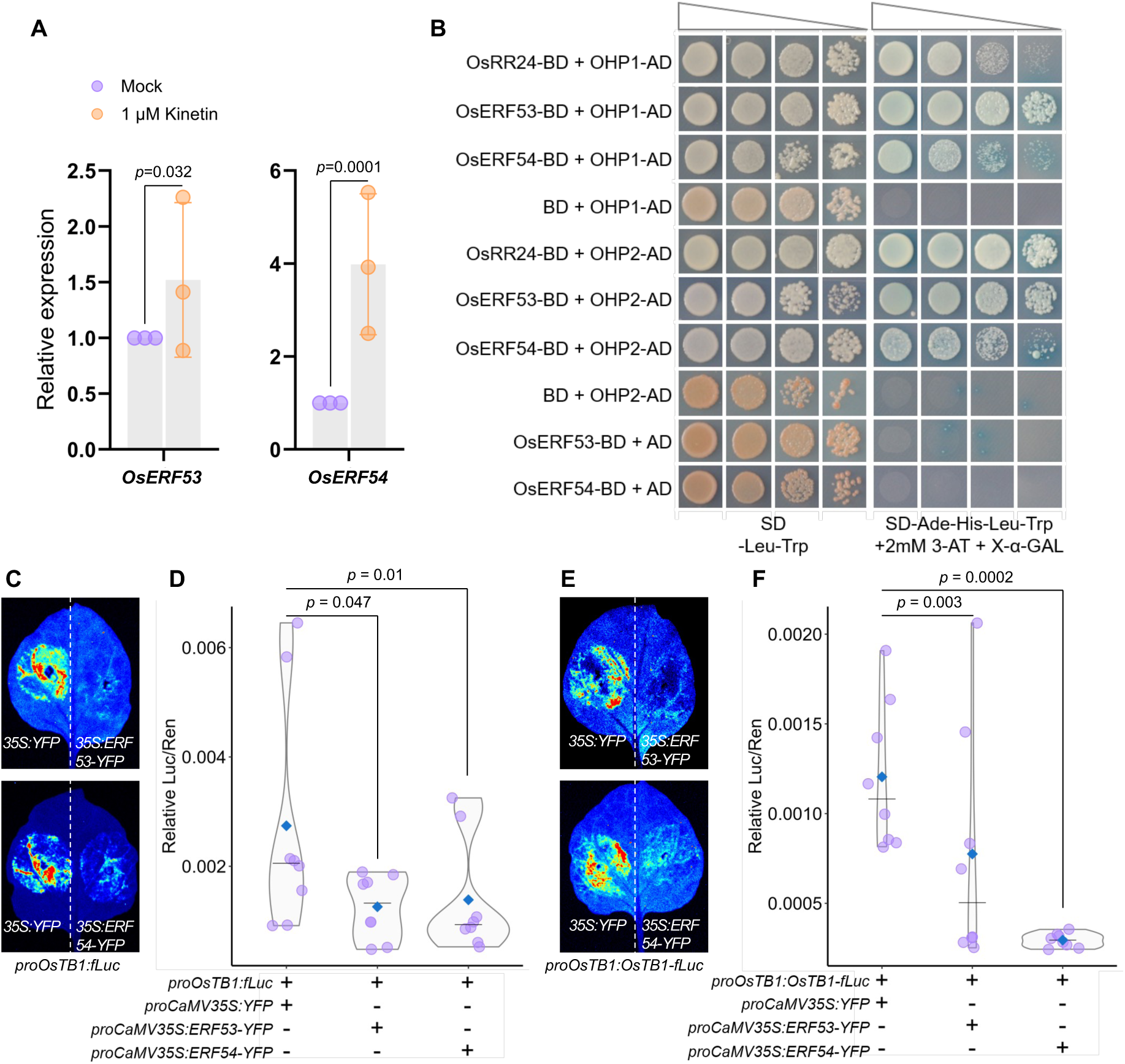
OsERF53/54 suppress *OsTB1* expression downstream of cytokinin. (A) Relative expression of *OsERF53* and *OsERF54* in minimal nitrogen media with and without 1μM Kinetin (n=3, p-values obtained from pairwise Student’s t test are indicated in the graphs). (B) Yeast two-hybrid assays showing an interaction between OsERF53/54 and OHP1/2. Transformed Y2H gold cells were grown on DDO (−Leu−Trp) and QDO (−Ade−Leu−Trp−His + 2 mM 3-aminotriazole [3AT] + 20 μg/ml X-α-Gal) media. Interactions between OsRR24 and OHP1/2 were used as positive controls, and combinations with empty vectors were used as negative controls. (C) Luciferase transactivation assay showing repression of *proOsTB1:fLuc* in *N. benthamiana* leaves co-infiltrated with control (*35S:YFP* in right panel) or TF (*35S:ERF53-YFP* or *35S:ERF54-YFP* in left panel). (D) Quantification of the LUC/REN ratio representing activity of the *OsTB1* promoter from (C). (E) Luciferase transactivation assay showing repression of *proOsTB1:OsTB1-fLuc* in *N. benthamiana* leaves co-infiltrated with control (*35S:YFP* in right panel) or TF (*35S:ERF53-YFP* or *35S:ERF54-YFP* in left panel). (F) Quantification of the LUC/REN ratio representing activity of the *OsTB1* translational fusion construct from (E). In (D) and (F), REN is used as an internal control. The blue diamonds and horizontal lines represent the mean and median values, respectively (n = 8; *p-*values obtained from pairwise Student’s t-test with control (*35S:YFP*) are indicated in the graphs).

### Cytokinin restricts polar auxin transport in tiller buds via *OsPIN1a* repression

Auxin signaling and polar auxin transport, together with cytokinin, emerged as key components in our transcriptomic analysis of nitrogen-induced transition from dormant to active tiller buds. Comparing nitrogen regimes and time points, we observed downregulation of *OsPIN1a* and most of the *PIN-LIKE* (*OsPILS*) transporters under nitrate– and ammonium-supplemented conditions (Fig. 5A), while other *OsPIN1* paralogs were upregulated. Given the higher transcript abundance of *OsPIN1a* and *OsPIN1b* in rice tiller buds, we focused further on these two transporters (Fig. S7A). Kinetin supplementation in minimal nitrogen consistently suppressed *OsPIN1a*, whereas *OsPIN1b* expression lacked any distinct pattern over 12 hours of exposure (Fig. S7B). We then supplemented CHX in combination with kinetin in minimal nitrogen to assess the role of canonical cytokinin signalling on *PIN* expression. *OsPIN1a* remained suppressed, while *OsPIN1b* showed slight upregulation, suggesting canonical cytokinin signalling-mediated regulation of *OsPIN1a* expression (Fig. 5B, S7C). Canonical cytokinin signalling, suppressing *OsPIN1a* downstream to nitrogen supply, further prompted us to check whether it is also regulated by OsERF53/54. The *OsPIN1a* promoter contained seven GCC-box motifs for ERF binding (Fig. S7D). Although the gene regulatory network predicted targeting of *OsPINa* only by OsERF53 (Fig. 2F), we tested the interaction of the *OsPIN1a* promoter (–2000 bp upstream of the start codon) with both ERFs in a transient dual-luciferase assay (Fig. S7E). Both OsERF53 and OsERF54 suppressed *OsPIN1a* promoter activity (Fig. 5C, D).

**Fig. 5.**
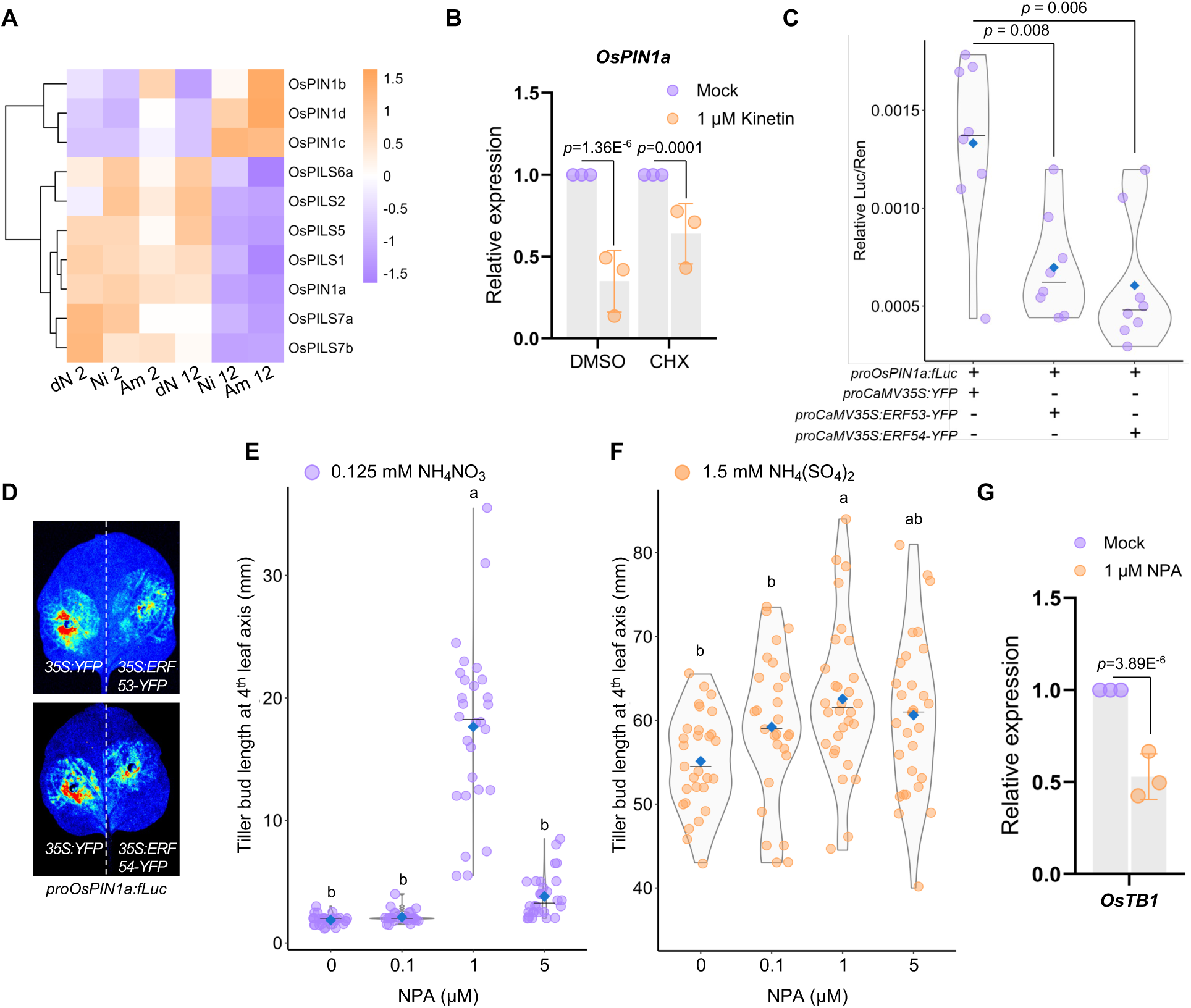
Cytokinin restricts polar auxin transport in tiller buds via *OsPIN1a* repression. (A) Heatmap representing the normalized expression values (z-score) of polar auxin transport genes in tiller buds at 2– and 12-hours across nitrogen regimes (dN/minimal nitrogen, Ni/sufficient nitrate, Am/sufficient ammonium). (B) Relative expression of *OsPIN1a* in minimal nitrogen media with and without 1μM Kinetin, either in mock (DMSO) or in 25 μM Cycloheximide (CHX) supplementation (n=3, *p*-values obtained from pairwise Student’s t test are indicated in the graphs). (C) Quantification of the LUC/REN ratio representing activity of the *OsPIN1a* promoter co-infiltrated with control (*35S:YFP*) or TF (*35S:ERF53-YFP* or *35S:ERF54-YFP*), where REN is used as an internal control. The blue diamond and horizontal line represent the mean and median values, respectively (n = 8; *p*-values obtained from pairwise Student’s t-test are indicated in the graphs). (D) Luciferase transactivation assay showing repression of *proOsPIN1a:fLuc* in *N. benthamiana* leaves co-infiltrated with control (*35S:YFP* in right panel) or TF (*35S:ERF53-YFP* or *35S:ERF54-YFP* in left panel). (E) & (F) Quantification of tiller bud length at the 4^th^ leaf axil of seedlings at four days post-supplementation of different concentrations of NPA in minimal nitrogen media (E) and in the presence of sufficient nitrogen (F). The blue diamonds and horizontal lines represent the mean and median values, respectively (n=27-30). The lowercase letters indicate significant differences based on one-way ANOVA and Tukey’s test (*p* < 0.05). (G) Relative expression of *OsTB1* in minimal nitrogen media with and without 1μM NPA (n=3, *p*-value obtained from pairwise Student’s t test is indicated in the graph). *OsGAPDH* was used as the endogenous control for qRT-PCR experiments.

Overall, the expression patterns of auxin transporters under nitrogen supplementation suggest that reduced polar auxin transport accompanies nitrogen-mediated tiller bud outgrowth. Since canonical cytokinin signaling represses *OsPIN1a*, and likely auxin transport, we tested whether pharmacological inhibition of auxin transport affects bud growth. The polar auxin transport inhibitor naphthylphthalamic acid (NPA) promoted tiller bud outgrowth in both nitrogen-deficient and ammonium-supplemented media (Fig. 5E, F, S8A, B), similar to cytokinin treatment. These results indicate that nitrogen and cytokinin suppress polar auxin transporters, promoting bud outgrowth, and that polar auxin transport acts as a negative regulator of this process. Consistent with this, NPA supplementation in minimal nitrogen also repressed *OsTB1* in tiller buds (Fig. 5G).

### Auxin induces *OsTB1* and *OsPIN1a* to inhibit bud outgrowth

Transcriptomic data strongly suggested antagonism between cytokinin and auxin signaling during nitrogen-triggered bud outgrowth (Fig. 2D, E), with most canonical auxin signaling and metabolism genes suppressed within 12 hours of nitrate or ammonium supply (Fig. S9). We therefore examined the role of auxin signaling, in addition to transport, in regulating bud outgrowth. Sufficient ammonium-containing media supplemented with increasing concentrations of picloram, a synthetic auxin analog, inhibited bud outgrowth in a dose-dependent manner (Fig. 6A, B). Progression of the cell cycle in tiller buds triggered by ammonium supply was also found to be impaired by picloram supplementation within 12 hours (Fig. 6C, D). These results highlight a repressive role of auxin in nitrogen-triggered tiller bud outgrowth.

**Fig. 6.**
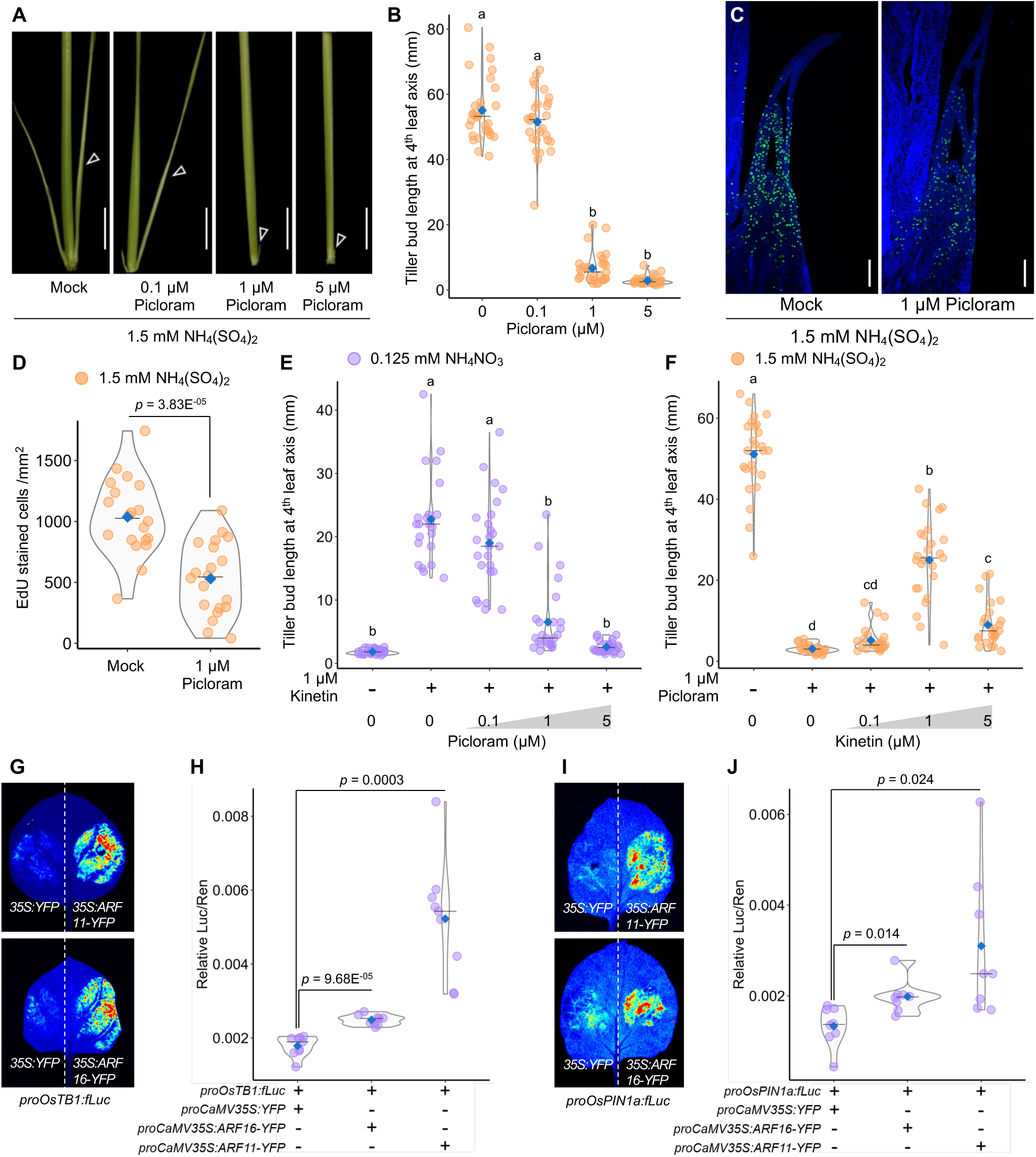
Auxin induces *OsTB1* and *OsPIN1a* to inhibit bud outgrowth. (A) & (B) Representative images (A, Scale = 1cm) and quantification (n=28-31, B) of tiller bud length at the 4th leaf axil of seedlings at four days post-supplementation with different concentrations of Picloram in sufficient nitrogen-containing media. (C) & (D) Representative images (C, Scale = 100 µm) and quantification (D) of EdU-stained cells in tiller buds at the 4^th^ leaf axil 12 hours after 1μM Picloram supplementation in sufficient nitrogen-containing media (n=15, *p*-value obtained from pairwise Student’s t test is indicated in the graph). (E) & (F) Quantification (n=23-25) of tiller bud length at the 4^th^ leaf axil of seedlings at four days post-supplementation in deficient nitrogen supplemented with 1μM Kinetin and different concentrations of Picloram (E) and in sufficient nitrogen supplemented with 1μM Picloram and different concentrations of Kinetin (F). (G) Luciferase transactivation assay showing repression of *proOsTB1:fLuc* in *N. benthamiana* leaves co-infiltrated with control (*35S:YFP* in right panel) or TF (*35S:ARF11-YFP* or *35S:ARF16-YFP* in left panel). (H) Quantification of the LUC/REN ratio representing activity of the *OsTB1* promoter from (G). (I) Luciferase transactivation assay showing repression of *proOsPIN1a:fLuc* in *N. benthamiana* leaves co-infiltrated with control (*35S:YFP* in right panel) or TF (*35S:ARF11-YFP* or *35S:ARF16-YFP* in left panel). (J) Quantification of the LUC/REN ratio representing activity of the *OsPIN1a* promoter from (I). In (H) and (J), REN is used as an internal control (n = 8; *p-*values obtained from pairwise Student’s t-test with control (*35S:YFP*) are indicated in the graphs). The blue diamonds and the horizontal lines in (B), (E), (F), (H), and (J) represent mean and median values, respectively. In (B), (E), and (F), the lowercase letters indicate significant differences based on one-way ANOVA and Tukey’s test (*p* < 0.05).

Having shown that auxin and cytokinin act antagonistically to control tiller bud outgrowth downstream of nitrogen, we next examined their interaction and combined effects with nitrogen supply. Under nitrogen-deficient conditions, kinetin was able to rescue tiller outgrowth. However, when picloram was added together with kinetin, it suppressed kinetin-induced bud outgrowth in a dose-dependent manner (Fig. 6E, S10A). In contrast, under sufficient ammonium supply, the inhibition caused by picloram was rescued by additional kinetin supplementation (Fig. 6F, S10B). These results suggest that auxin and cytokinin attenuate each other’s effects on tiller outgrowth without fully overriding the other. Based on these pharmacological assays, we hypothesized that both hormones might converge on a common target to regulate nitrogen-dependent tiller outgrowth.

Since cytokinin represses *OsTB1* and *OsPIN1a*, both inducers of tiller bud dormancy, we tested whether auxin also regulates their expression. Promoter scans revealed four and ten AuxRE motifs in *OsTB1* and *OsPIN1a*, respectively (∼2 kb upstream of the start codon), which are essential for Auxin Response Factor (ARF) binding (Fig. S11A, B) (Rienstra *et al*., 2023). Gene regulatory networks indicated that both genes are targeted by multiple ARFs, hub and non-hub alike (Fig. S5B, C). Among them, we selected OsARF11 and OsARF16 for further study, as both are clade-A ARFs with genetic evidence for negative regulation of rice tiller outgrowth (Matthes *et al*., 2019; Uzair *et al*., 2021). Transient dual-luciferase assays showed that OsARF11 and OsARF16 significantly induced *OsTB1* and *OsPIN1a* promoter activity (Fig. 6G, H, I, J). These findings demonstrate that auxin signaling directly activates *OsTB1* and *OsPIN1a* transcription via OsARF11/16, promoting tiller bud dormancy. We propose that under sufficient nitrogen, reduced auxin signaling in tiller buds lowers *OsTB1* and *OsPIN1a* expression, facilitating bud outgrowth. In summary, auxin and cytokinin, through OsARF11/16 and OsERF53/54, respectively, converge on shared targets such as *OsTB1* and *OsPIN1a*, exerting opposing effects on their expression.

### *OsERF53/54* promotes nitrogen-triggered tiller bud outgrowth

Based on our results, we propose that the auxin–cytokinin antagonism serves as a key regulatory module, integrating nitrogen signals to determine tiller bud fate. The results connected the previously isolated nodes of tiller bud outgrowth, such as *OsTB1*, *OsPIN1a*, *OsARF11*, and *OsARF16*, in the nitrogen-triggered tiller outgrowth network and assembled a more coherent view of the underlying regulatory framework governing this process. Moreover, we identified two key components, *OsERF53* and *OsERF54*, that were not known for their involvement in tiller outgrowth and lack genetic evidence substantiating their involvement in the process. To this end, we examined natural diversity in rice accessions to determine whether sequence variations in these genes are associated with tillering capacity. Using genetically diverse rice accessions (Supplementary Table S1), we assessed tiller numbers under sufficient ammonium supply and found significant variation among these accessions (Fig. 7A). We then analyzed publicly available sequence data of the selected rice varieties for sequence variation in *OsERF53/54* (∼2 kb upstream of the start codon to the 3′ UTR) and identified multiple haplotypes for each gene (Supplementary dataset S5) (Mansueto *et al*., 2017; Wang *et al*., 2018a). We selected three haplotypes comprising most accessions per gene from these and compared average tiller numbers (Fig. S12A, B). Accessions carrying *OsERF53* Hap12 showed significantly higher tiller numbers than Hap5, with multiple SNPs in both the promoter region and gene body (Fig. S12C). Hap9 was nearly identical to Hap12, differing only by a 6 bp promoter insertion, and displayed a similar phenotype (Fig. S12A, C). However, since Hap9 included only three accessions, we excluded it from further experiments. *OsERF54* Hap4 exhibited higher tillering than other haplotypes and carried a unique 15 bp CDS deletion (Fig. S12B, D). We classified the haplotypes into two groups for subsequent analysis. The Hap A group included *OsERF53* Hap12 and *OsERF54* Hap4, mostly high-tillering accessions. In contrast, the Hap B group included *OsERF53* Hap5 and *OsERF54* Hap6 and Hap10, mostly low-tillering accessions (Fig. S12C, D). Many of the selected rice accessions harbored Hap A of both genes (Fig. S12E). Similarly, many other accessions had Hap B of both genes (Fig. S12F). Rice accessions with Hap A of both genes had significantly higher average tiller numbers than those with Hap B of both genes (Fig. 7B, S12E, F).

**Fig. 7.**
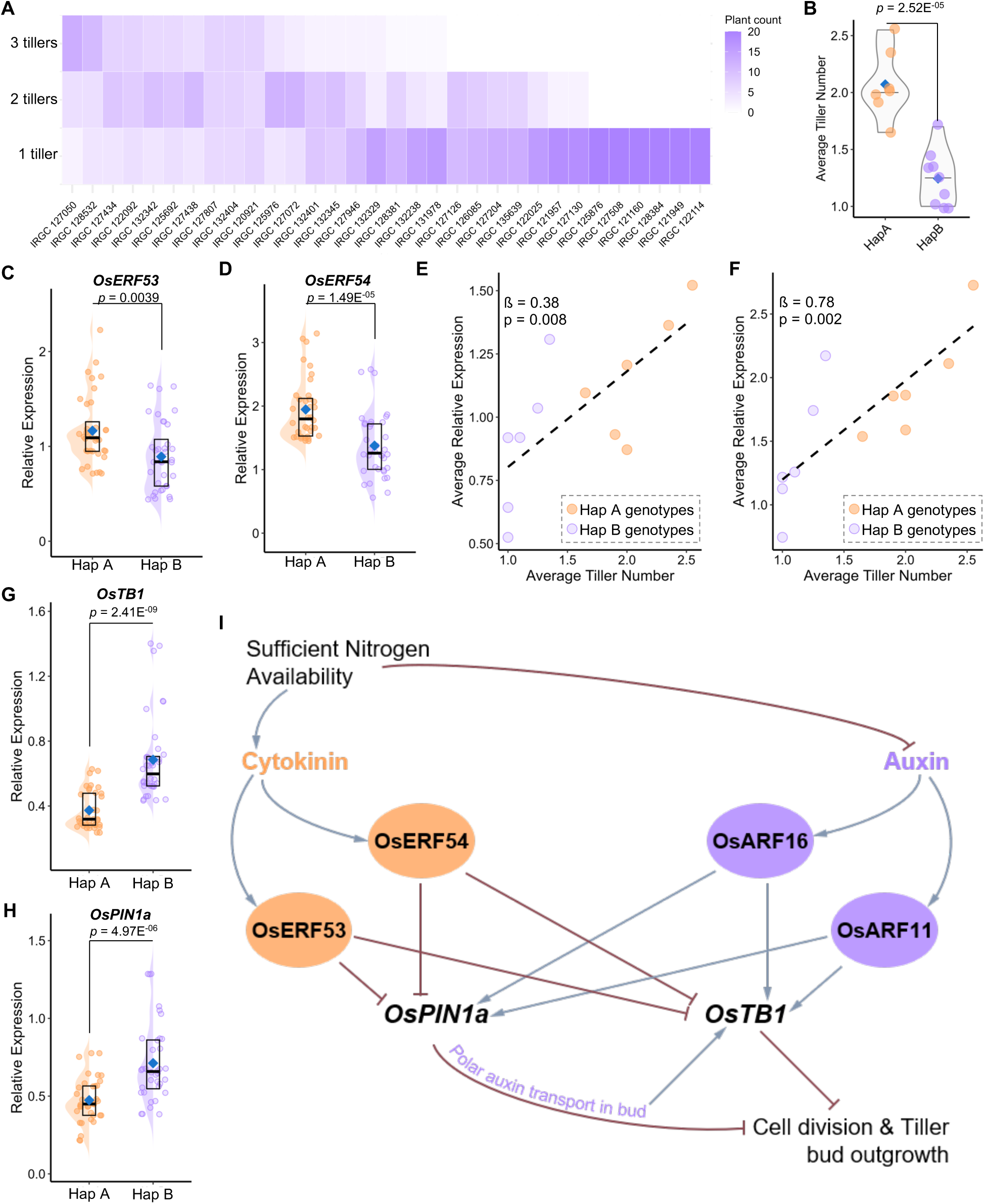
OsERF53/54 promotes nitrogen-triggered tiller bud outgrowth. (A) Quantification of tiller number (main culm + tiller buds>10mm in length) at 4 weeks after growing in sufficient ammonium-containing media. The heatmap represents the number of plants from each genotype having either 1, 2, or 3 tillers. (B) Average tiller number in genotypes having shared HapA or HapB for both *OsERF53* and *OsERF54*. The *p*-value indicated in the graph is based on a pairwise Student’s t-test. (C) & (D) Relative expression of *OsERF53* (C) and *OsERF54* (D) in response to sufficient ammonium at 6 hours in comparison to minimal nitrogen. The blue diamonds represent mean values, and boxes represent median and interquartile ranges (n=6, 2 biological and 3 respective technical replicates for 6 varieties for both HapA and HapB, *p*-values obtained from pairwise Student’s t test are indicated in the graphs). (E) & (F) Scatterplots representing the average expression of *OsERF53* (in E) and *OsERF54* (in F) in selected genotypes in the y-axis plotted against the average tiller number obtained from (A) in the x-axis. The dashed lines represent the linear regression line (ß = slope). *p*-values are indicated in the graphs. (G) & (H) Relative expression of *OsTB1* (G) and *OsPIN1a* (H) in response to sufficient ammonium at 6 hours in comparison to minimal nitrogen. The blue diamonds represent mean values, and boxes represent median and interquartile ranges (n=6, 2 biological and 3 respective technical replicates for 6 varieties for both HapA and HapB, *p*-values obtained from pairwise Student’s t test are indicated in the graphs). (I) A schematic representation of nitrogen-triggered tiller bud outgrowth in rice. Adequate nitrogen activates cytokinin signaling and suppresses auxin signaling in tiller buds. Nitrogen and cytokinin both induce the transcription of *OsERF53/54*, which in turn represses *OsTB1* and *OsPIN1a*. Conversely, auxin, through OsARF11/16, promotes the expression of these genes. Repression of *OsPIN1a* limits polar auxin transport in tiller buds, which further reduces *OsTB1* expression. Lower *OsTB1* activity releases buds from dormancy, allowing cell division and subsequent outgrowth.

We then selected six representative accessions from Hap A and Hap B to assess whether genomic variation correlated with expression patterns. Seedlings were grown under minimal nitrogen until the 5th leaf was fully expanded, then half were supplemented with sufficient ammonium. Gene expression from the 4th leaf axil tiller buds was compared between ammonium-supplemented and minimal nitrogen plants. *OsERF53* expression was significantly higher in Hap A accessions, with a positive correlation to the average tiller number of corresponding accessions (Fig. 7C, E). *OsERF54* showed a similar trend with an even stronger correlation (Fig. 7D, F). Consistent with their predicted regulatory role, *OsTB1* and *OsPIN1a* were more strongly repressed by ammonium in Hap A accessions with more tiller numbers than Hap B accessions with lower tiller numbers (Fig. 7G, H). Together, these results indicate that natural genetic variation in rice influences the expression differences in *OsERF53/54*, which regulate tillering under sufficient nitrogen supply. Higher ERF expression was correlated with higher tiller production, supporting *OsERF53/54* as positive regulators in nitrogen-triggered tiller outgrowth.

## Discussion

Tiller bud outgrowth in rice is a critical determinant of yield, bridging genetic potential and realized performance under fluctuating environments (Takai, 2024). Environmental constraints such as nitrogen availability strongly influence tiller outgrowth, often overriding genetic potential (Luo *et al*., 2020; Takai, 2024). While genetic regulators of tiller outgrowth have been extensively identified through screens of high– and low-tillering rice mutants, their integration with environmental cues remains poorly understood (Arite *et al*., 2009; Hao *et al*., 2009; Minakuchi *et al*., 2010; Duan *et al*., 2019; Fang *et al*., 2020). Notably, the understanding of the integration of these cues during the early transition from bud dormancy to active growth is limited due to the lack of methods to precisely trigger tiller activation in rice. In dicots such as Arabidopsis and pea, early stages of axillary bud outgrowth are routinely studied using decapitation to remove apical dominance (Cao *et al*., 2023; Beveridge *et al*., 2023; Nahas *et al*., 2024). This provides a well-established system to dissect early regulatory steps of bud activation. However, similar methods are not suitable for cereals like rice due to the weak apical dominance and the distinct positioning of the shoot apical meristem (Kebrom *et al*., 2013; Xia *et al*., 2020). Due to this limitation, studying the early regulatory steps of rice tillering remains challenging. We manipulated nitrogen supply, which triggered bud activation in a controlled manner, precisely synchronizing the transition from dormancy to growth (Xu *et al*., 2015). This approach enabled us to explore the early stages of nitrogen-induced tiller outgrowth. Using IR64 indica rice seedlings, we profiled transcriptomic changes under contrasting nitrogen conditions. Our findings highlight cytokinin–auxin antagonism as a central module acting downstream of nitrogen signaling to regulate early bud outgrowth.

Rice tiller buds remain dormant when nitrogen is limiting, immediately triggering outgrowth in seedlings with sufficient nitrogen supplementation (Fig. 1B, C). Of the two major nitrogen forms in agricultural soils, we found ammonium (NH₄⁺) to be more effective than nitrate (NO₃⁻) in promoting outgrowth (Fig. 1B–E). This aligns with earlier studies reporting higher tiller and panicle numbers in rice supplied with NH₄⁺ as the sole nitrogen source (Luo *et al*., 2017; Yi *et al*., 2019a; Hou *et al*., 2021b). Rice’s preference for NH₄⁺ reflects its adaptation to submerged, anaerobic conditions (Gu & Yang, 2022). Unlike NH₄⁺, NO₃⁻ uptake shows an initial lag in hydroponic culture before increasing (Sasakawa & Yamamoto, 1978). Coupled with the higher energy demand of NO₃⁻ assimilation, this lag likely delays its effect on tillering (Bloom, 2015). Our transcriptomic analysis revealed nitrogen form-specific early responses. Within two hours, NH₄⁺ induced substantial transcriptional changes, while NO₃⁻ had little effect (Fig. 2A, B). By 12 hours, however, responses to both nitrogen forms overlapped significantly (Fig. 2A, B). Cytokinin-related responses were enriched after 2 hours of NH₄⁺ supply (Fig. S2A). Cytokinin biosynthesis in response to nitrogen is known to depend on glutamine, a product of nitrogen assimilation, rather than on the specific nitrogen form (Kamada-Nobusada *et al*., 2013). Notably, indica cultivars carry a more efficient OsNR2 allele, conferring higher nitrate reductase activity and faster NO₃⁻ assimilation (Gao *et al*., 2019). Thus, the slow uptake but rapid assimilation of NO₃⁻ in indica likely explains the expression dynamics observed in NO₃⁻-fed tiller buds.

Among phytohormone pathways, cytokinin was the only one enriched in upregulated genes, while several others were enriched in downregulated genes (Fig. S2A, B). Dormant buds showed enrichment of stress– and dormancy-related hormones such as ABA, consistent with its known role in bud dormancy in maize and Arabidopsis (González-Grandío *et al*., 2013; Dong *et al*., 2019; van Es *et al*., 2024). Among growth hormones, GA and strigolactone were enriched in downregulated terms, both known suppressors of tillering (Arite *et al*., 2009; Hao *et al*., 2009; Minakuchi *et al*., 2010; Umehara *et al*., 2010; Fang *et al*., 2020; Wu *et al*., 2020). Interestingly, auxin signaling and transport were also enriched in downregulated terms, but have not yet been integrated into the molecular framework underlying tiller outgrowth in rice. Co-expression modules revealed a near-bimodal expression pattern: the negatively correlated turquoise and blue modules together contained over 90% of DEGs (Fig. S3A, S4B). This likely reflects the large overlap between upregulated, as well as downregulated, genes at 12 hours under both nitrogen forms (Fig. 2B). Notably, the 2-hour NH₄⁺ response also overlapped substantially with the later 12-hour response for both upregulated and downregulated genes and comprised a few unique genes (Fig. S2C, D). Auxin– and cytokinin-related terms were separated into these two antagonistic modules, and shared more hub genes than other hormones (Fig. 2D, S4B). This recurrent inverted pairing prompted us to examine their roles more closely, given that auxin-cytokinin antagonism regulates many developmental processes, including shoot branching in Arabidopsis (Müller & Leyser, 2011; Müller *et al*., 2015; Kotov & Kotova, 2023).

Our results confirm that cytokinin promotes tiller bud outgrowth downstream of nitrogen supply by inducing cell division within 12 hours (Fig. 3B–E). This is consistent with enhanced tillering in transgenic rice lines with increased cytokinin biosynthesis, stability, or transport (Yeh *et al*., 2015; Zhao *et al*., 2019). Cytokinin suppressed *OsTB1*, a central inhibitor of tiller outgrowth, establishing it as a primary cytokinin-responsive gene (Fig. 3F). However, canonical cytokinin-related TFs such as type-B Response Regulators (BRRs) were clustered in the blue module, which was associated with dormancy (Fig. S3C-E). Genetic evidence also suggests *OsBRR*s have little effect on tiller number, with increases seen only in higher-order mutants (Worthen *et al*., 2019; Yamburenko *et al*., 2020; Li *et al*., 2022a). Instead, two Cytokinin Response Factor (CRF) family members, *OsERF53/54*, emerged as potential positive regulators. Their expression patterns, role in nitrogen-induced gene regulatory networks (Fig. 2E, F, S6A), and ability to directly repress *OsTB1* promoter activity (Fig. 4C-F) link nitrogen-triggered enhanced cytokinin signaling to tiller bud outgrowth. In Arabidopsis, CRFs are known to alter shoot architecture in response to nitrogen availability (Abualia *et al*., 2022). Supporting this, variation in the *OsERF53/54* genomic region across several rice accessions was associated with altered tillering capacity, and their expression correlated positively with ammonium-induced bud outgrowth (Fig. 7C–F, S12), establishing them as positive regulators of tiller outgrowth downstream to adequate nitrogen supply.

Most polar auxin transport genes were downregulated after 12 hours of NH₄⁺ or NO₃⁻ supply or kinetin treatment (Fig. 5A, S5A, B). Alongside *OsTB1*, cytokinin suppressed *OsPIN1a* via OsERF53/54 (Fig. 5B-C, S5B, F). In Arabidopsis, auxin canalization via AtPIN1 is essential for lateral bud outgrowth (Prusinkiewicz *et al*., 2009), while *AtPIN3*, *PIN4*, and *PIN7* also promote branching downstream of cytokinin (Bennett *et al*., 2006; Waldie & Leyser, 2018). Similarly, CsPIN3-mediated auxin canalization regulates cucumber branching (Shen *et al*., 2019). These findings support a triggering role for auxin transport during axillary bud outgrowth in dicots. However, auxin transport appears to function differently in rice. The rice genome encodes 12 PIN genes with distinct tissue-specific expression patterns and roles in tillering (Wang *et al*., 2009; Yan *et al*., 2023). *OsPIN9*, a poaceae-specific transporter, promotes auxin transport and tillering under NH₄⁺ supply (Hou *et al*., 2021a). Although *OsPIN9* expression was reported in tiller vasculature, we did not detect *OsPIN9* transcripts in dormant or active buds during early outgrowth. *OsPIN2*, another positive regulator (Chen *et al*., 2012), was also absent. In contrast, knockdown or knockout of *OsPIN1/OsPIN1b* increases tiller number, and higher-order mutants of *OsPIN1a*, *PIN1c*, and *PIN1d* show enhanced tillering (Xu *et al*., 2005; Li *et al*., 2019, 2022b; Wang *et al*., 2022). Intriguingly, our data also showed *OsPIN1* paralogs as the major *PIN*s expressed in buds (Fig. 5A), and genetic evidence suggests that they suppress bud outgrowth. These findings suggest auxin canalization from rice tiller buds is unlikely during early outgrowth stages. Supporting this, bud outgrowth was induced by auxin transport inhibitors NPA and TIBA in the monocot orchid *Cremastra appendiculata*, comparable to decapitation (Lv *et al*., 2018), highlighting divergence in auxin-mediated branching between monocots and dicots. Consistently, our pharmacological assays showed that polar auxin transport restricts tiller bud activation under both nitrogen-deficient and sufficient conditions (Fig. 5D, E, S8A, B). Previous studies reported similar increased tillering in rice supplemented with NPA at specific concentrations (Hao *et al*., 2009; Hou *et al*., 2021a). Unlike Arabidopsis, where branching depends on release from apical dominance and axillary buds compete to become auxin sources (Bennett *et al*., 2006), weaker apical dominance in rice (Xia *et al*., 2020) may underlie these distinct roles of auxin transport during tiller bud outgrowth.

In contrast to cytokinin, rice transgenics with reduced auxin sensitivity or enhanced inhibition of auxin signaling show increased tillering (Xia *et al*., 2012; Jin *et al*., 2016). Consistent with this, we observed that auxin suppressed ammonium-induced bud cell division and tiller outgrowth (Fig. 6A–D). Curiously, auxin and cytokinin could not fully negate each other’s effects when supplied together, regardless of nitrogen status (Fig. 6E, F, S10A, B). This suggests that rather than acting hierarchically, auxin and cytokinin converge on common targets to regulate outgrowth. Supporting the negative influence of auxin on tiller outgrowth, overexpression of an auxin-resistant OsIAA10 protein increased tiller number, whereas *iaa10* mutants had fewer tillers (Jin *et al*., 2016; Qin *et al*., 2020). OsIAA10 interacts with auxin-responsive transcription factors OsARF11, OsARF12, and OsARF16, which may act as regulators of tiller outgrowth (Qin *et al*., 2020). While *OsARF12* knockout and overexpression altered tiller angle (Qi *et al*., 2012; Wang *et al*., 2014; Zhao *et al*., 2023), its role in outgrowth remains unclear. In contrast, *arf11* and *arf16* knockout lines showed high tillering, identifying them as negative regulators (Uzair *et al*., 2021). Although OsARF11 and OsARF16 were not identified as hub transcription factors in our contrasting co-expression modules, they were deeply connected in the gene regulatory network underlying nitrogen-triggered tiller bud outgrowth. Both OsARF11 and OsARF16 transiently activated *OsTB1* and *OsPIN1a* promoters (Fig. 6G–J), corroborating our hypothesis that these genes act as convergent targets of auxin and cytokinin signaling. Our placement of *OsTB1*, *OsPIN1a*, *OsARF11*, and *OsARF16* in the nitrogen-triggered tiller outgrowth network is supported by prior genetic evidence for their individual roles in tillering (Minakuchi *et al*., 2010; Li *et al*., 2019; Uzair *et al*., 2021). We connected these previously fragmented nodes into a hierarchical network and incorporated cytokinin-driven OsERF53/54, providing a comprehensive regulatory module beyond previously characterized components of rice tillering.

In conclusion, we synchronized tiller bud outgrowth in rice seedlings by manipulating nitrogen supply and analyzed the transcriptional changes underlying the early stages of bud activation under sufficient nitrogen. Among the regulatory pathways involved, we focused on cytokinin–auxin antagonism, which converges on common targets to determine bud fate. We found that cytokinin promotes bud outgrowth by repressing *OsTB1* and *OsPIN1a* through transcriptional regulation by OsERF53/54 (Fig. 7I). In contrast, auxin maintains dormancy by inducing these genes via OsARF11/16. Sufficient nitrogen supply promotes cytokinin signaling in tiller buds and suppresses auxin transport and signaling, collectively tipping the balance toward bud activation (Fig. 7I). Our study integrates established regulators with newly identified factors, presenting a unified framework of the nitrogen-triggered bud outgrowth network. Finally, we detected the natural variation in the genomic sequences of *OsERF53/54* that influences the responsiveness of rice accessions to nitrogen, highlighting a potential resource for breeding rice cultivars with improved nitrogen use efficiency.

## Acknowledgments

This work was supported by the core funding from the National Institute of Plant Genome Research and Rice Network Project (BT/Ag/Network/Rice/2019–20) from the Department of Biotechnology, Ministry of Science and Technology, India. SC acknowledges CSIR-JRF and AD acknowledges UGC-JRF, respectively. We thank NIPGR Central Instrumentation Facility (CIF) and Plant Growth Facility for their support.

## Author contributions

S.C.-Investigation, data curation, visualization, formal analysis, validation, and writing-original draft, A.D.-Formal analysis, visualization, and writing-review and editing. A.K.S. – Conceptualization, funding acquisition, and writing-review and editing. A.R.-Conceptualization, supervision, funding acquisition, formal analysis, and writing-review and editing. All authors have read and approved the final version of the manuscript.

## Data Availability Statement

The data that support the findings of this study are available in the supplementary material of this article. The quality-filtered reads, which were used to get the normalized read counts and for differential gene expression analysis, are deposited in the NCBI Short Read Archive under accessions SRR35297547-SRR35297552.

## Supporting information

### Supplemental Datasets

**Supplemental Dataset S1.** List of statistically significant differentially expressed genes for each pair-wise comparison.

**Supplemental Dataset S2.** List of statistically significant gene ontology categories (biological processes) enriched for differentially expressed genes and hub genes of co-expression modules.

**Supplemental Dataset S3.** List of genes and hub genes categorized in different co-expression modules.

**Supplemental Dataset S4.** List of canonical genes belonging to cell-cycle, tillering, and phytohormone signaling pathways from rice, curated from literature.

**Supplementary Dataset S5.** Genomic variation, haplotype combinations, and expression variation of *OsERF53* and *OsERF54* in different rice accessions.

### Supplementary Figures

**Fig. S1** Temporal expression dynamics of bud outgrowth marker genes.

**Fig. S2** Biological processes enriched in nitrogen-responsive DEGs.

**Fig. S3** Co-expression modules associated with nitrogen regimes.

**Fig. S4** Hormone-related pathways enriched in co-expression modules.

**Fig. S5** Targets of cytokinin– and auxin-related canonical transcription factors.

**Fig. S6** Expression of *ERF*s and binding sites in *OsTB1*.

**Fig. S7** Repression of polar auxin transporters during bud emergence.

**Fig. S8** Reduced auxin transport promotes bud outgrowth.

**Fig. S9** Auxin metabolism and signaling under nitrogen regimes.

**Fig. S10** Combinatorial effects of nitrogen, cytokinin, and auxin on bud outgrowth.

**Fig. S11** AuxRE distribution in *OsTB1* and *OsPIN1a* promoters.

**Fig. S12** Haplotype variation in *OsERF53/54* genomic sequences.

### Supplementary Tables

**Supplementary Table S1.** Rice accessions used in this study from the IRRI 3K panel.

**Supplementary Table S2.** Composition, stock, and the final concentrations of nutrients for modified Yoshida media.

**Supplementary Table S3.** Primers used in this study.

**Supplementary Table S4**: R packages used in this study.

