## Supplemental Figures S1 – S12 for "A cytokinin-auxin antagonistic module regulates nitrogen-triggered tiller outgrowth in rice"

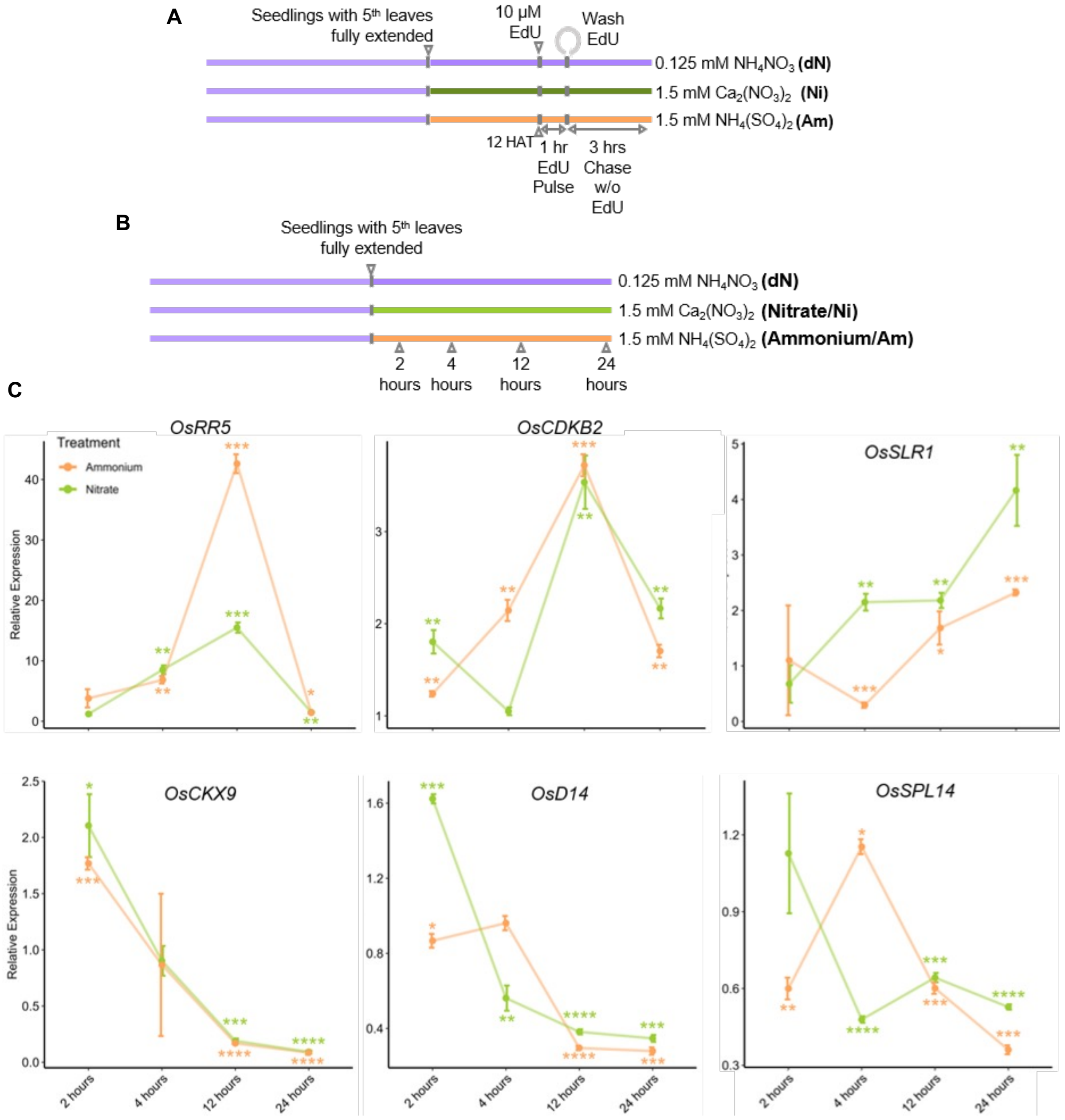


**Fig. S1 Temporal expression dynamics of bud outgrowth marker genes.** (A) Schematic diagram showing the experimental conditions for the cell-proliferation assay. Seedlings grown on minimal nitrogen-containing Yoshida media were continued in minimal nitrogen (dN) or supplemented with sufficient nitrate (Ni) or ammonium-containing media (Am). After 12 hours, 10 µM EdU was supplemented in the respective media for an hour (Pulse), and then washed. Tiller buds were kept in the respective media after washing for an additional 3 hours (Chase) before fixation. (B) Schematic diagram showing the experimental conditions for gene expression analysis. Seedlings grown on minimal nitrogen-containing Yoshida media were continued in minimal nitrogen (dN) or supplemented with sufficient nitrate (Ni) or ammonium-containing media (Am). Tiller buds were harvested at the mentioned time points for expression analysis. (C) Relative fold change of marker genes in Ni- or Am-treated buds compared to dN-treated buds at different time points. Data shown are mean ± SD, stars indicate significant differences based on Student's t-test ****$p$<0.0001, ***$p$< 0.001, **$p$ < 0.01, *$p$< 0.05.

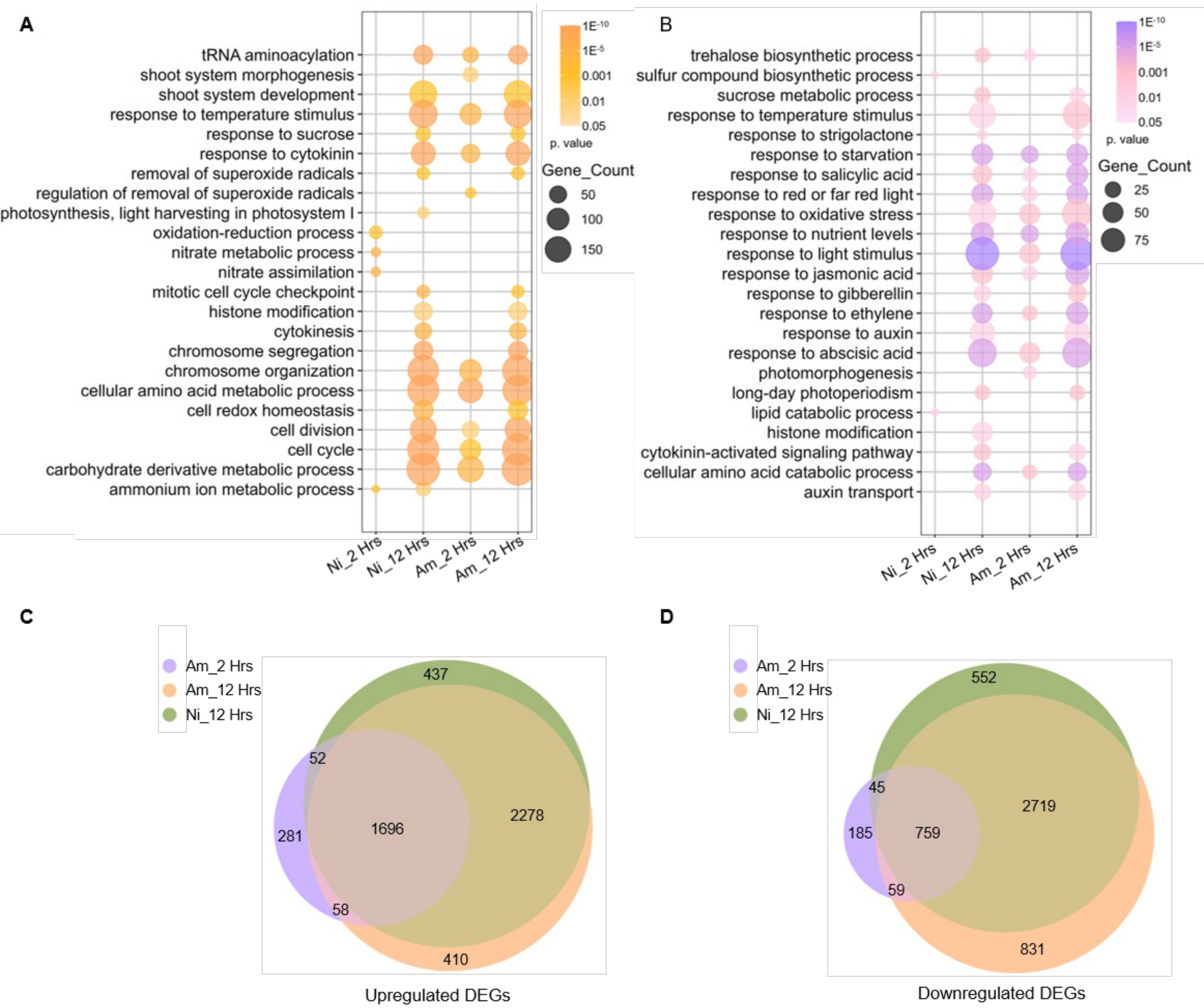


**Fig. S2 Biological processes enriched in nitrogen-responsive DEGs.** (A) & (B) Bubble plot displaying representative GO terms (biological process) enriched for the Upregulated (A) and Downregulated genes (B) in Ni- and Am-treated seedlings compared to dN-treated seedlings at respective time points. (C) & (D) Euler diagrams illustrating the numbers and overlap of Upregulated (C) and Downregulated (D) DEGs in Am and Ni supplementation in comparison to dN at respective time points. dN - minimal nitrogen, Ni – sufficient nitrate, Am – sufficient ammonium.

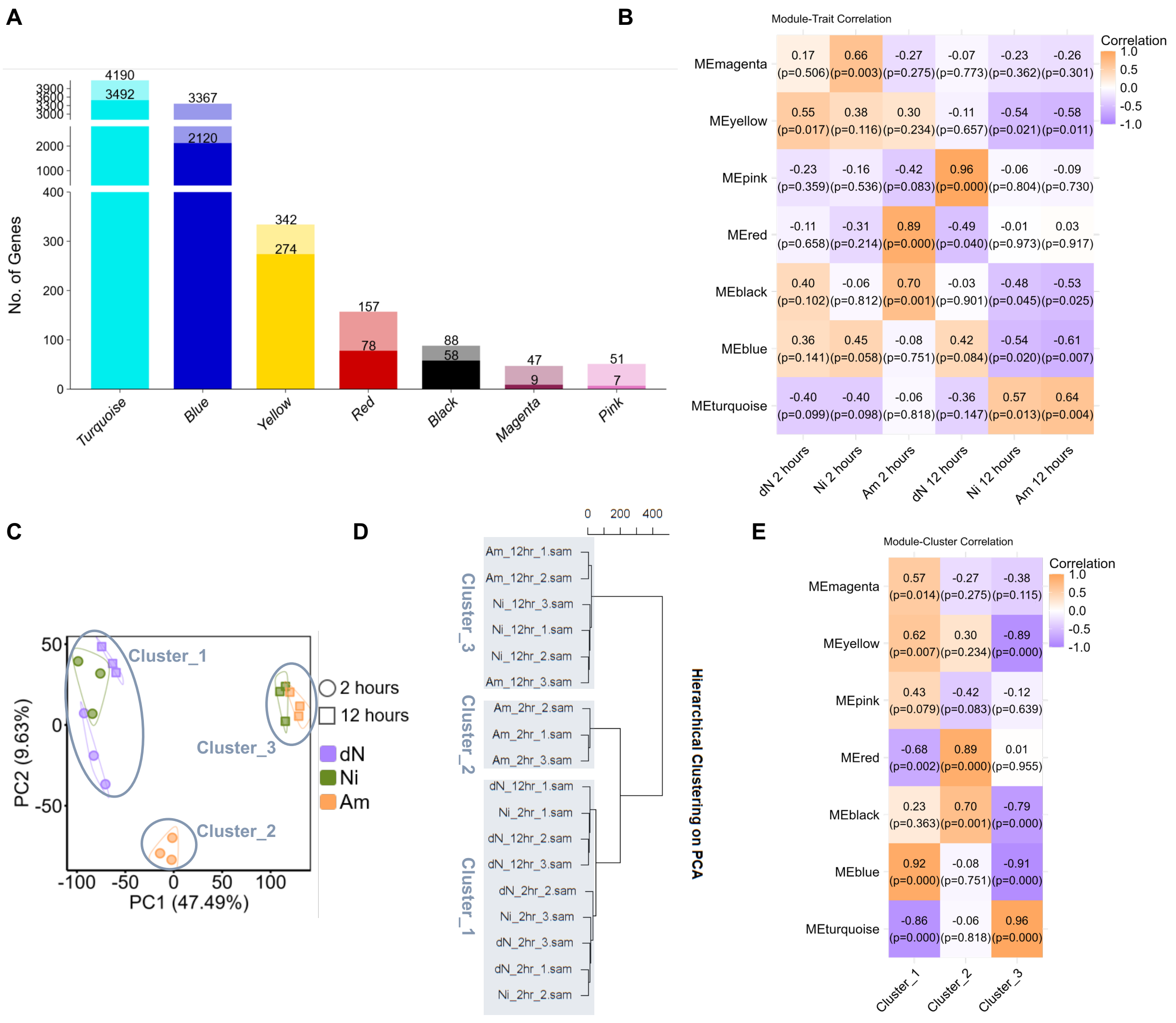


**Fig. S3 Co-expression modules associated with nitrogen regimes.** (A) Stacked bars representing the total number of genes in each module. The lower, darker parts represent the number of hub genes in each module. (B) Module-sample correlation derived from WGCNA analysis. The row corresponds to modules, and the columns correspond to samples. Module names are mentioned on the left. The color gradient of the cells represents the correlation coefficient (values mentioned in cells) of the corresponding module and sample. The significance of the association is denoted by the p-value (shown in parentheses). dN - minimal nitrogen, Ni – sufficient nitrate, Am – sufficient ammonium. (C) & (D) Principal-component analysis (PCA, C) and Hierarchical Clustering on PCA (D) of gene expression profile obtained from Fig. 2A. 18 samples were split into three distinct sample clusters through Hierarchical clustering. (E) Module-cluster correlation derived from WGCNA analysis. The row corresponds to modules, and the columns correspond to sample clusters. Module names are mentioned on the left. The color gradient of the cells represents the correlation coefficient (values mentioned in cells) of the corresponding module and cluster. The significance of the association is denoted by the p-value (shown in parentheses).

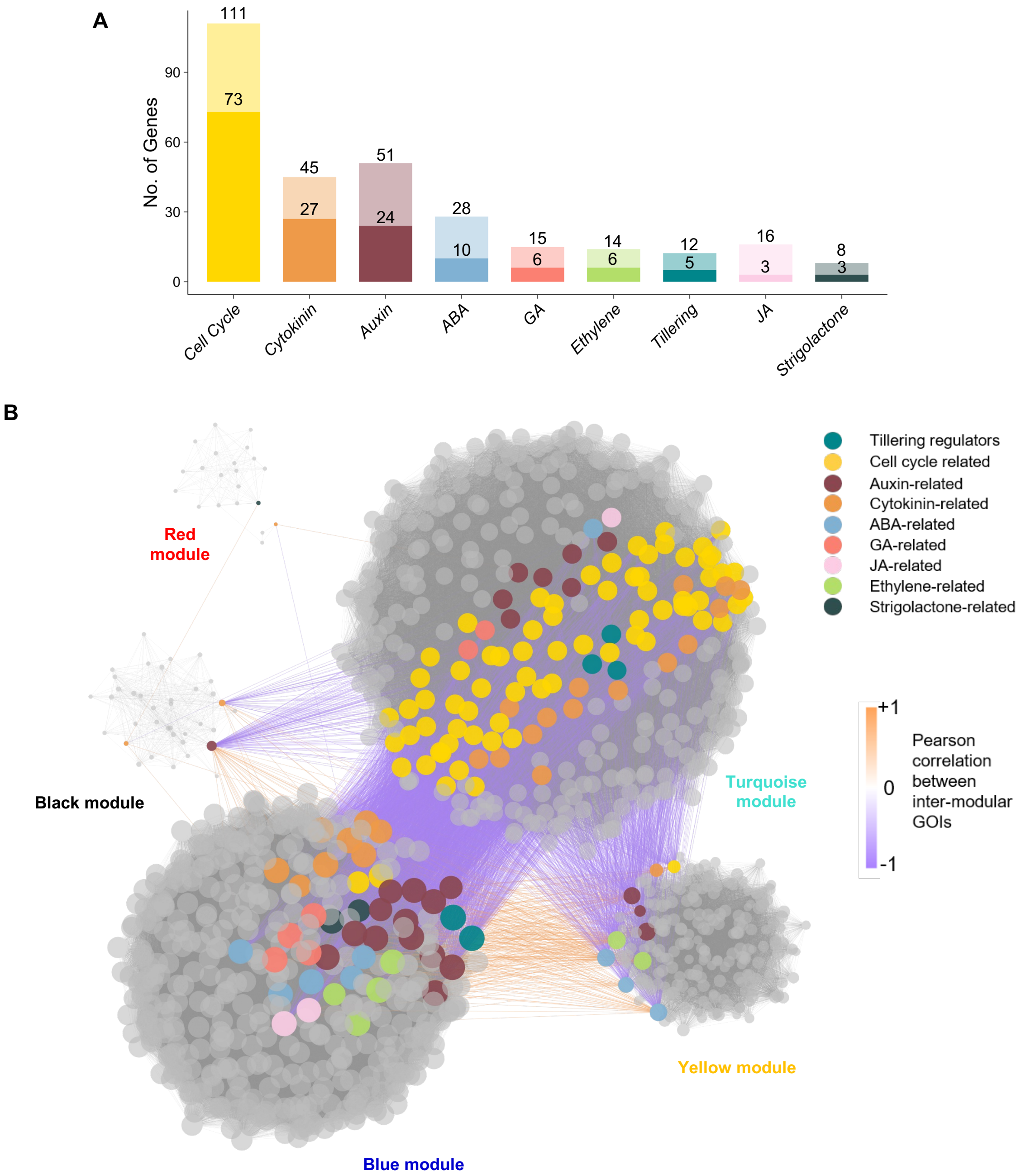


**Fig. S4 Hormone-related pathways enriched in co-expression modules.** (A) Stacked bars representing the total number of expressed canonical genes of different pathways of interest in hub genes across all modules. The lower, darker parts represent the number of hub genes in each pathway belonging to any module. (B) Co-expression network of hub GOIs and their immediate neighbors (Up to 50) across modules. Node size represents connectivity within the module. Colored edges represent Pearson correlations between intra-modular GOIs.

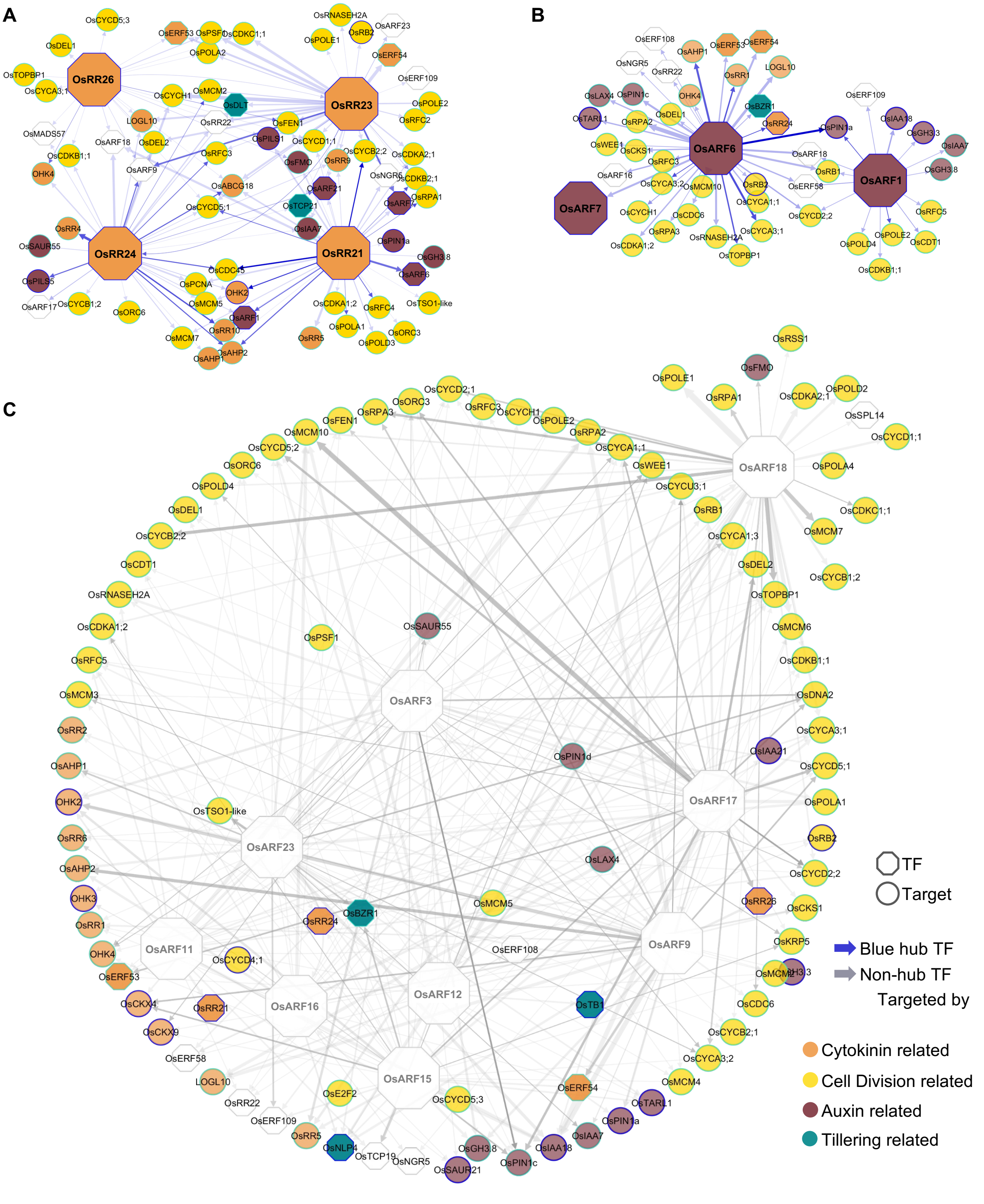


**Fig. S5 Targets of cytokinin- and auxin-related canonical transcription factors.** Gene regulatory networks for cell cycle, auxin, cytokinin, and tillering-related genes from the turquoise and blue modules targeted by (A) Hub BRRs, (B) Hub ARFs, and (C) Non-hub ARFs. Colors of node borders depict module assignment, and edge colors represent the module of regulatory TF. Edge thickness represents the number of binding sites from −2kb to +1kb from the TSS of target genes. and Edge transparency represents Edge weight. Unfilled TF nodes represent TFs from auxin or cytokinin pathways that are not hub-TFs from the blue or turquoise modules.

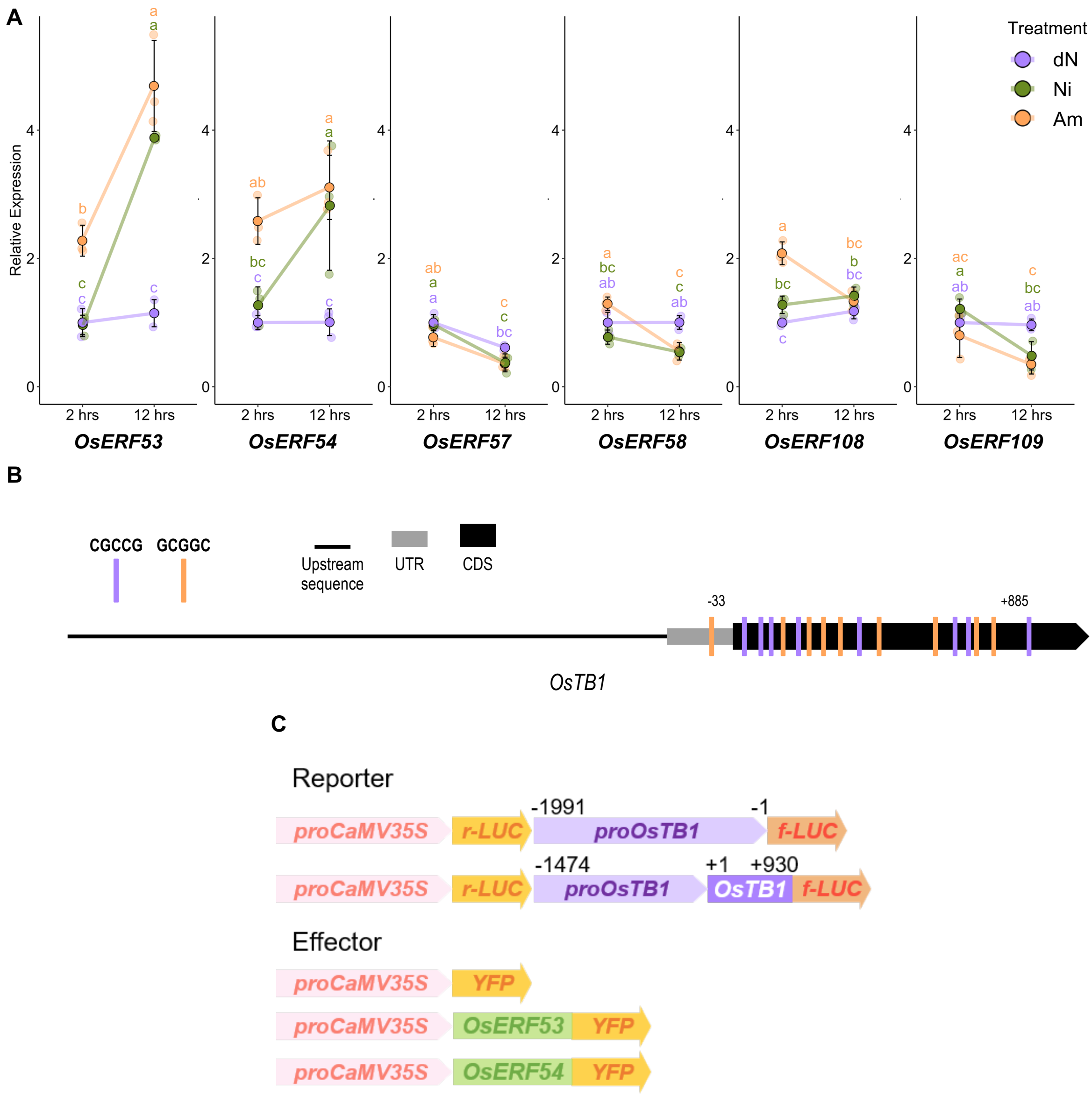


**Fig. S6 Expression of *ERF*s and binding sites in *OsTB1*.** (A) Normalized expression values of *OsERF* genes (belonging to the *OsCRF* family) scaled against 2 hours dN samples. The letters indicate significant differences based on one-way ANOVA and Tukey's test ($P < 0.05$). The color of the letters corresponds to the treatments. dN - minimal nitrogen, Ni – sufficient nitrate, Am – sufficient ammonium. (B) Schematic representation of the *OsTB1* gene and upstream sequence for the presence of putative ERF binding motifs (positions of motifs most distant from the start codon (+1) are mentioned on either upstream or downstream). (C) Schematic representation of the constructs used in the transient transactivation assays (ATG start codon was considered as +1).

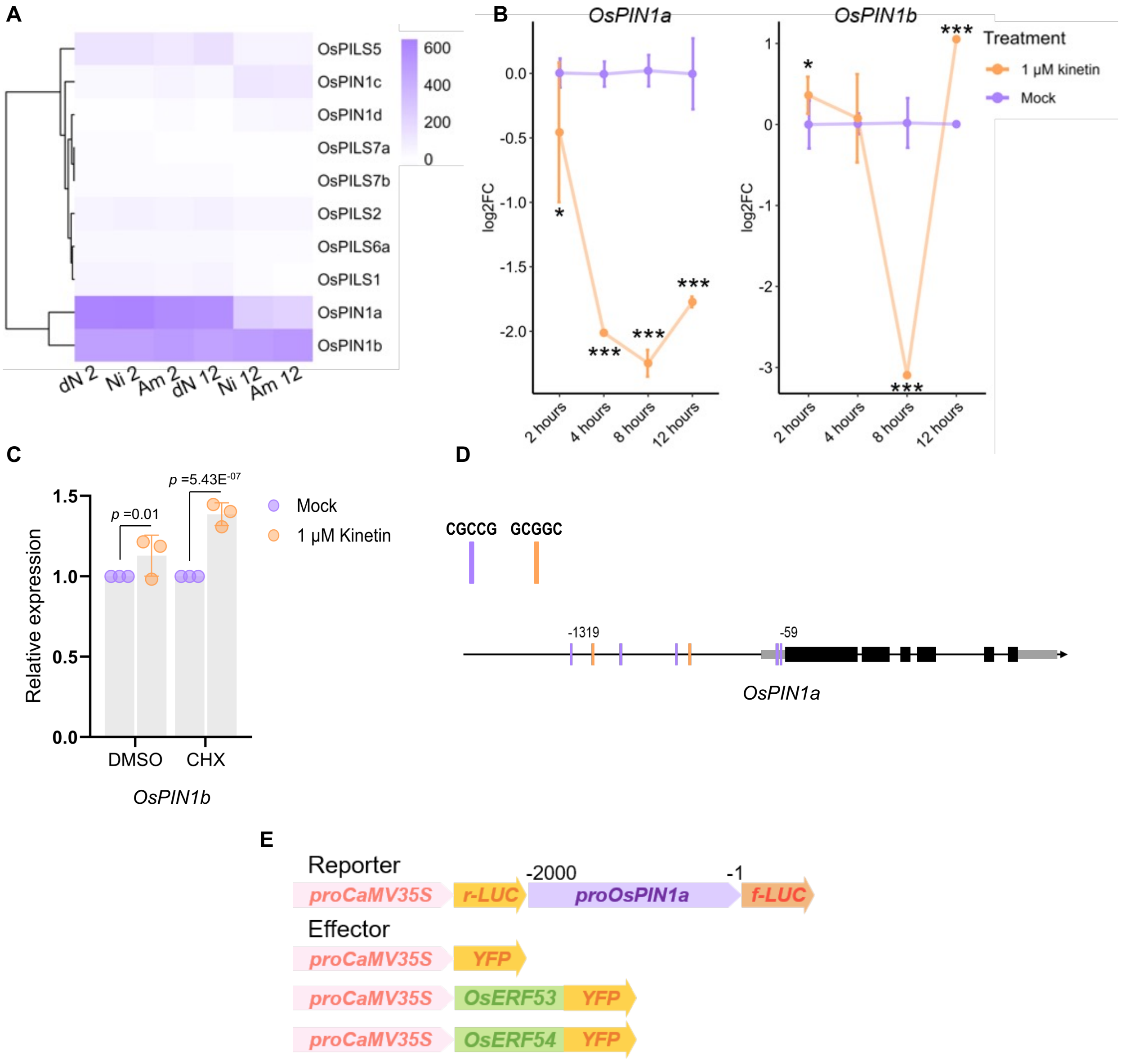


**Fig. S7 Repression of polar auxin transporters during bud emergence.** (A) Heatmap representing the absolute normalized counts of polar auxin transport genes in tiller buds at 2- and 12-hours across nitrogen regimes (dN/minimal nitrogen, Ni/sufficient nitrate, Am/sufficient ammonium). (B) Relative fold change in *OsPIN1a* and *OsPIN1b* in 1 µM kinetin supplementation compared to mock in minimal nitrogen media. Data shown are mean ± SD. Stars indicate significance based on Student's t-test (***$p$< 0.001, **$p$ < 0.01, *$p$< 0.05). (C) Relative expression of *OsPIN1b* in minimal nitrogen media with and without 1µM Kinetin, either in mock (DMSO) or in 25 µM Cycloheximide (CHX) supplementation (n=3, $p$-values obtained from pairwise Student's t test are indicated in the graphs). (D) Schematic representation of the *OsPIN1a* gene and upstream sequence for the presence of putative ERF binding motifs (nearest and farthest positions of motifs upstream of the start codon (+1) are mentioned). (E) Schematic representation of the constructs used in the transient transactivation assays (ATG start codon was considered as +1).

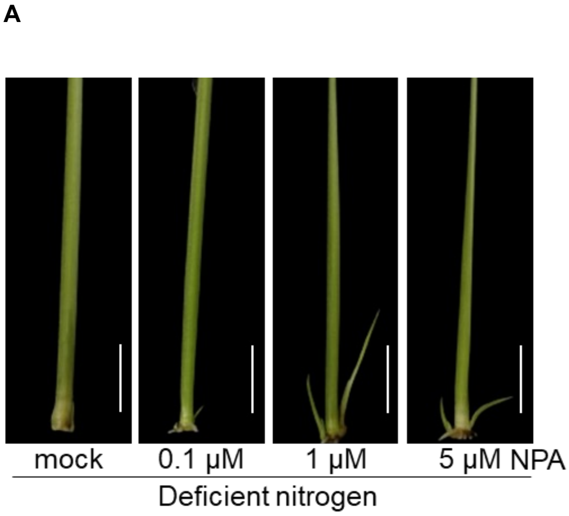


**Fig. S8 Reduced auxin transport promotes bud outgrowth.** (A) & (B) Representative images of tiller bud length at the 4th leaf axil of seedlings at four days post-supplementation of different concentrations of NPA in deficient nitrogen media (A) and in the presence of sufficient nitrogen (B).

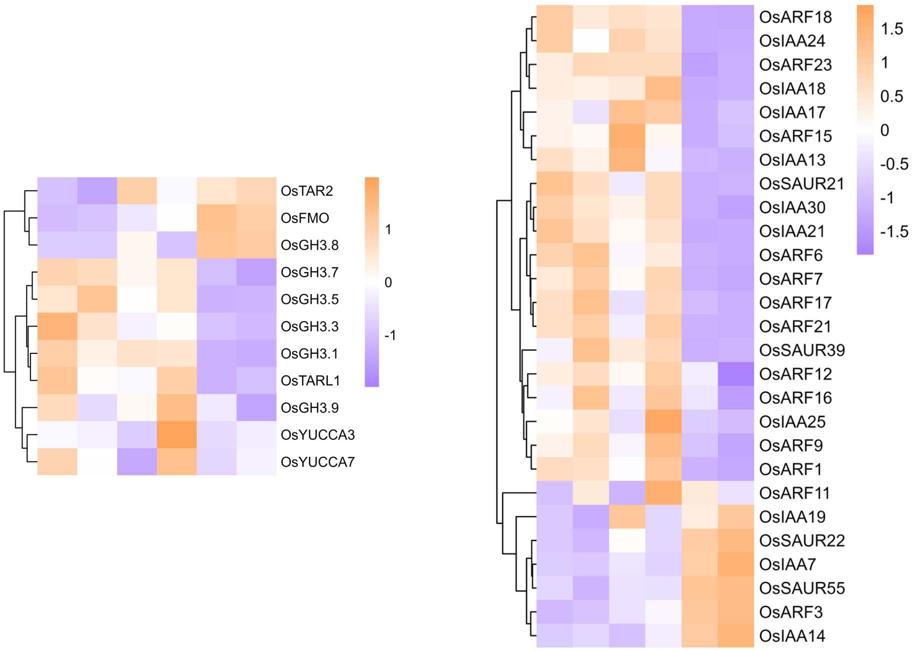


**Fig. S9 Auxin metabolism and signaling under nitrogen regimes.** Heatmap representing the normalized expression values (z-score) of auxin metabolism and signaling genes in tiller buds at 2- and 12-hours across nitrogen regimes (dN/minimal nitrogen, Ni/sufficient nitrate, Am/sufficient ammonium).

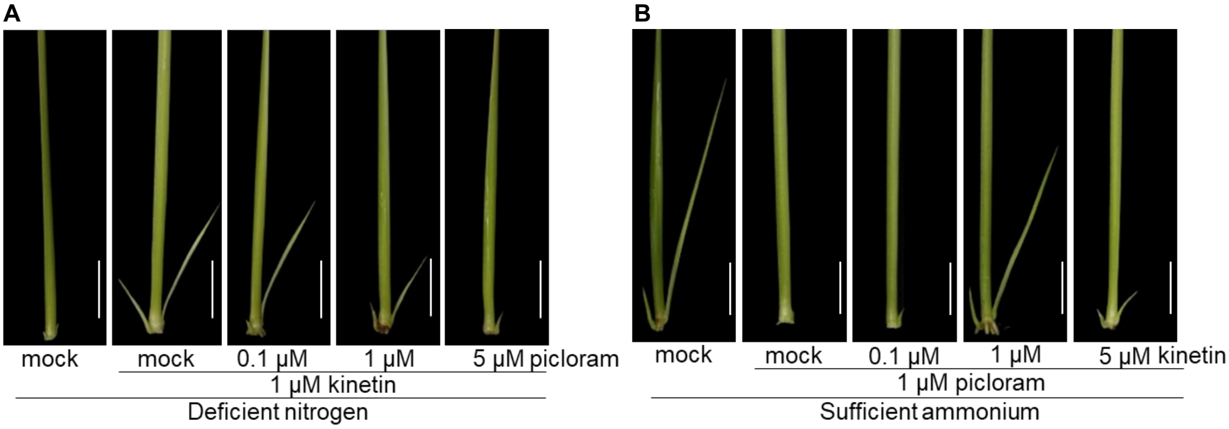


**Fig. S10 Combinatorial effects of nitrogen, cytokinin, and auxin on bud outgrowth.** (A) & (B) Representative images of tiller bud length at the 4th leaf axil of seedlings at four days post-supplementation in deficient nitrogen supplemented with 1µM Kinetin and different concentrations of Picloram (A) and in sufficient nitrogen supplemented with 1µM Picloram and different concentrations of Kinetin (B).

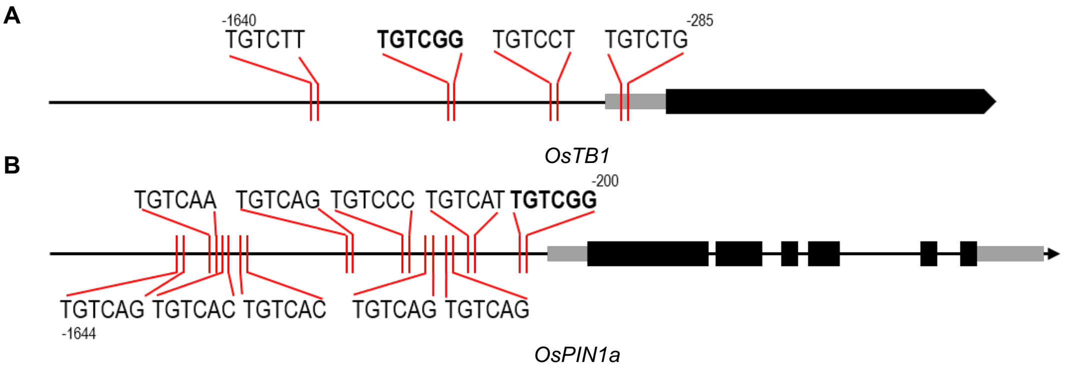


**Fig. S11 AuxRE distribution in *OsTB1* and *OsPIN1a* promoters.** Schematic representation of the *OsTB1* (A) and *OsPIN1a* (B) gene and upstream sequence for the presence of canonical (represented in bold) and non-canonical AuxREs (nearest and farthest positions of motifs upstream of ATG are mentioned).

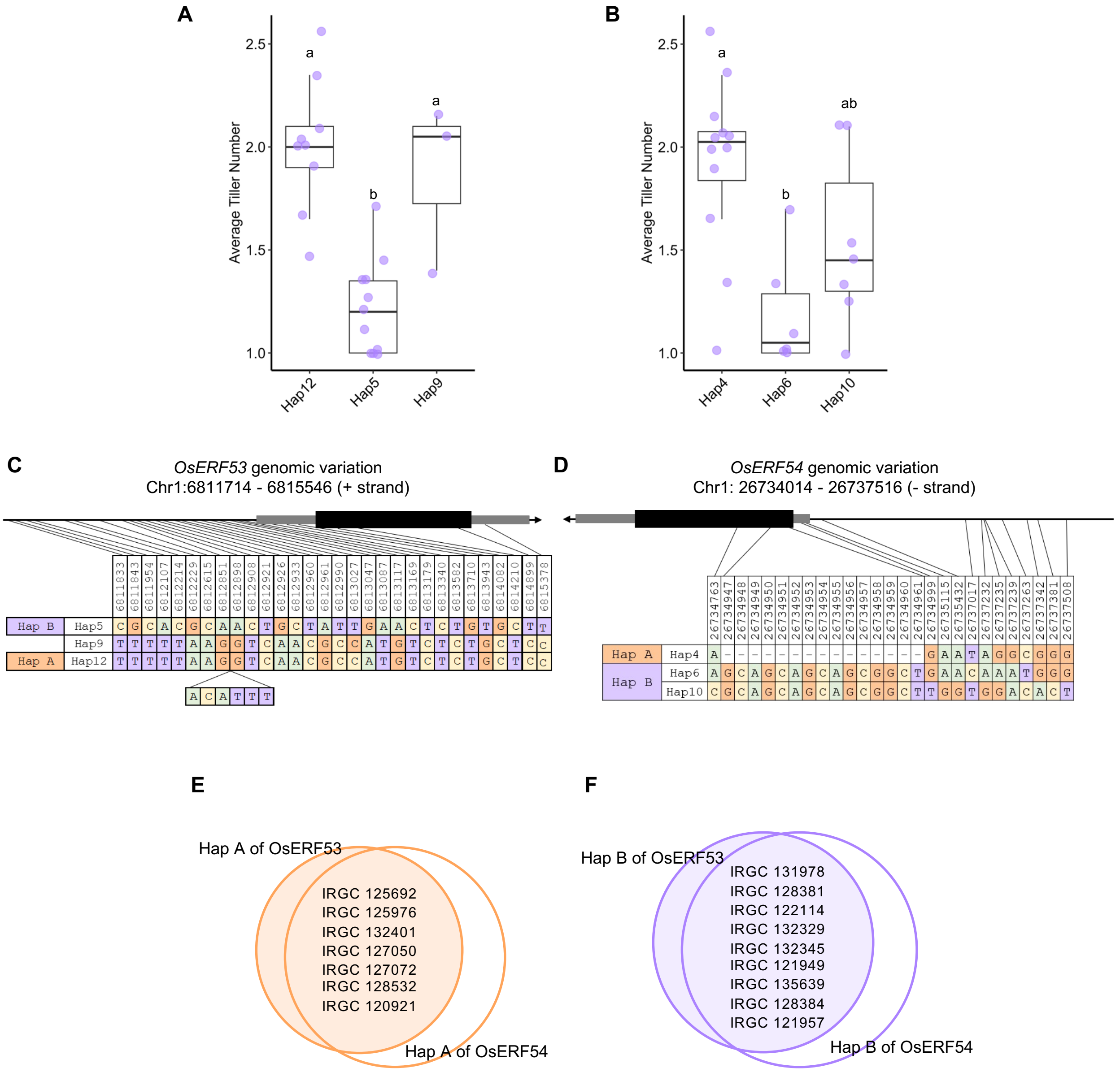


**Fig. S12 Haplotype variation in *OsERF53/54* genomic sequences.** (A) & (B) Average tiller number in the varieties of the most abundant haplotype for *OsERF53* (A) and *OsERF54* (B). The lowercase letters indicate significant differences based on one-way ANOVA and Tukey's test ($p < 0.05$). Data for (A) and (B) is obtained from Fig7A. Boxplots show medians, interquartile ranges, and min-max values with individual data points superimposed as colored dots. (C) & (D) Sequence variations in the selected haplotypes of *OsERF53* (*C*) and *OsERF54* (*D*) loci. (E) & (F) Common genotypes belonging to HapA (E) and HapB (F) of *OsERF53* and *OsERF54.*
